# Exploring media representation of the exotic pet trade: taxonomic, framing, and language biases in peer-reviewed publications and newspaper articles

**DOI:** 10.1101/2024.05.01.592090

**Authors:** Jon Bielby, Gail E. Austen, Kirsten M. McMillan, Shannen M. Wafflart

## Abstract

1. The exotic pet trade is a global industry with considerable implications for a range of taxa and stakeholders. The scale of the trade means it receives coverage in both popular and scientific media, and some narratives may receive more attention than others. As these media play an important role in shaping public opinion, policy, and legislation, we should consider and acknowledge biases and language use when reporting on the exotic pet trade.
2. We use 320 peer-reviewed journal articles, and 191 newspaper articles on the exotic pet trade between 2001 and 2020 to investigate the frequency of use, citation rate, and language-use across framing categories and taxonomic foci within and between media-types.
3. Our results suggest consistent biases in reporting of the trade within and between media-types, highlighting limitations in both. Aspects of welfare were underrepresented in peer-reviewed articles relative to other framings, but it was the most common focus of newspaper articles.
4. If the exotic pet trade is to develop into a more ethical, sustainable, economically beneficial sector, then reassessing our narratives, improving knowledge flow, and encouraging interdisciplinary and comprehensive debates within the field will be essential parts of the process.

## Introduction

The global pet trade in non-domesticated companion animals (hereafter exotics; Alves 2012; De la Fuente et al. 2023) is vast. It is estimated to be worth billions of US dollars per annum, comprising of millions of individuals, across a broad range of taxonomic groups (Watters et al. 2022), and supply chains of varying size and legality (Bush et al. 2014). Recent analyses suggest that between 1999 and 2015, over 187 million individual vertebrates were imported into the USA for the pet trade market (Sinclair et al. 2021). However, the demand for exotic pets, as part of the international wildlife trade, is extremely difficult to quantify accurately (Hughes 2021). Therefore, these figures are likely to vary considerably between region and taxonomic group, but one thing is clear: both the trade and its repercussions are considerable.

Given its size and scope, the impact of this trade is felt at a range of scales from the individual animal to the ecosystem (Hughes 2021). Advocates of the trade highlight how it can be economically and culturally important within range states (Alves et al. 2012, Alves & Rocha 2018) and how the trade may provide sustainable livelihoods to communities within some regions (Robinson et al. 2018a, King 2019). It can also enrich the lives of pet owners (Haddon et al. 2021; De la Fuente et al. 2023), and advances in techniques and knowledge sharing regarding husbandry and nutrition have been beneficial to conservation efforts supporting *in situ* populations (Pasmans et al. 2017). On the flip side, detrimental effects of the pet trade include individuals suffering reduced welfare (Pasmans et al. 2017); population decline of source animals (Tingley et al. 2017; Nijman et al. 2018; Mandimbihasina et al. 2020); spread of infectious disease (Fitzpatrick et al. 2018); and invasive species causing significant damage to native species and ecosystems (Lockwood et al. 2019). The diverse costs and benefits outlined above cause it to be a controversial subject that continues to generate political and scientific debate.

Interpreting, framing, and communicating the trade in exotics is subjective, yet incredibly important (Challender et al. 2021, Natusch et al. 2021a, Easter et al. 2023). The complex costs and benefits associated with it provide opportunity for studies to be communicated from various angles. In this article we refer to these as different framings (see also Baker et al. 2013), which is to say that certain aspects of the subject may be highlighted to be more memorable or salient to the reader (Entman 1993). The use of different framings may be highly dependent upon the interests and beliefs of the author(s) (Natusch et al. 2021a), the target audience, and the objective of the research (Moorhouse et al. 2017). For example, philosophical biases of the author(s) may lead to inadvertent misrepresentation of data (see Natusch et al. 2021a; but see Edwards et al. 2021); open-access databases can be used to misrepresent the level of threat, trade-related policies and the nature of trade itself (Challender et al. 2021); and peer-reviewed literature and popular media may exhibit systematic taxonomic biases in acknowledging or reporting issues in wildlife and its trade (Baker et al. 2013; Feber et al. 2017). Similarly, the motivations of the author(s) and funder(s) can affect how data are framed, with the objective of influencing stakeholder or consumer choices (Moorhouse et al. 2017). Increasingly, as the pet trade and related policy come under scrutiny in several countries (e.g., CITES Secretariat 2022, Scottish Animal Welfare Commission 2022), peer-reviewed papers, grey literature, and popular media play a key role in shaping legislation via both public opinion and direct reporting to government bodies. All media types help form our perceived reality (Entman 1993), shaping public perception and attitudes surrounding the urgency and magnitude of such issues (e.g., Walker et al. 2019; Hammond et al. 2022). Consequently, we need to consider and understand framing use, consciously or otherwise, within communications on the exotic pet trade and how this varies across taxonomic groups and media types.

The subject of framing within wildlife trade has previously been discussed. A decade ago, Baker et al. (2013) published an overview of a range of types of literature relevant to the more general area of ‘wildlife trade’ (rather than the exotic pet trade, as in this article), looking to identify the presence of different aspects of welfare. They found that most articles were centred around conservation (71%), whereas only 17% of papers contained references to animal welfare, such as referring to a particular element of the five domains model (Mellor 2016). The welfare of animals within any trade or utilisation scheme should be an important factor for all stakeholders involved, from free-market advocates to those seeking a ban on the activity. Further, being concerned with the constituent individuals being traded is relevant, regardless of their source, provenance, domestic status, or conservation needs. Therefore, a specific focus of our analyses is to identify how the use of welfare as a framing of research has changed in the decade since Baker et al. published their broader findings; despite the large size of the trade and the risks it poses, it is unclear how the peer-reviewed literature has kept pace and whether the relative lack of research on the subject has been addressed. This is important, because the presence of biases in the framing of wildlife trade literature suggests that knowledge-gaps will exist, limiting the evidence-base with which legislation and policies may be informed.

In this article we aim to provide an overview of how the exotic pet trade is communicated in both the peer-reviewed literature and in one format of popular media – newspaper articles - with a specific focus on the use of welfare as a framing tool and its prevalence in both types of literature. First, we determine which framings are most widely used and compare their frequency both within and between media types. Second, we used natural language processing techniques to summarise how language use varied between the two media-types, specific to the different framings. Additionally, for peer-reviewed literature we also highlight taxonomic biases and describe how those foci covary with the framing category used and investigate whether those two factors cause variation in the citation rate of a peer-reviewed paper. By analysing and evaluating these two media types in this way, this article seeks to provide an overview of the biases and knowledge gaps in the reporting of the exotic pet trade. Only by identifying and filling these gaps and balancing these systemic biases will we be able to provide a more comprehensive knowledge base for evidence informed policy on the trade.

## Methods

Ethical statement: As our data collection and analyses were focused on publicly available peer-reviewed and newspaper articles no ethical approval was deemed necessary.

### Collating articles and defining frames and taxonomic focus: Peer-reviewed literature

To systematically identify and collate peer-reviewed publications we used specifically developed search terms in Scopus and Web of Science databases limiting the search to years from 2001 to 2021 inclusive. We focused on articles published in English because, while non-English language peer-reviewed papers are available and becoming more common (Amano et al. 2021), during this period English was the most consistently used language in scientific publishing (and with this came associated problems; Amano et al. 2023). Full details of the search terms and processes followed are outlined in Table S1 and S2 and Fig S1. Throughout the data collection process, PRISMA (Preferred Reporting Items for Systematic Review and Meta-Analyses) (Page et al. 2020) was used as a guide, to provide clarity in our methods and ensure replicability. A list of the details of resulting peer-review articles will be available to download from the LJMU data repository.

We defined peer-reviewed publications and their framing largely following Baker et al. (2013) and via preliminary investigation of the publications yielded by the literature search. The Baker paper outlined the high prevalence of conservation, human health, and economic framings, and was focussed on identifying welfare focussed content. We therefore included these four as framing categories, broadening the human health category to include the infectious disease risk posed by the pet trade to native species. Additionally, we added an ‘invasive species’ framing category, as this is a large ecological cost associated with the global trade in exotics (Lockwood et al. 2019). This process therefore resulted in five non-mutually exclusive framing categories: ‘Conservation’, ‘Disease’, ‘Economics’, ‘Invasive Species’, and ‘Welfare’. To allocate peer-reviewed publications into framing categories we developed a list of search terms (Table S3), the presence of which in the title, abstract and keywords was used to allocate articles to categories. If a publication fit into more than one framing category the paper was allocated to a separate ‘Multiple’ grouping, and if no criteria were met it was allocated to ‘No/other frame’. For peer-reviewed literature individual keyword searches were used to identify the taxonomic focus (S4 Table). A ‘Multiple’ grouping was used for articles with multiple taxonomic foci, and an ‘Other’ category if none of the search terms were met.

### Collating articles and defining frames: Newspaper articles

The search terms ti(‘exotic pet*’) AND ti(‘trade’) were used to interrogate the ProQuest database for entries of English language newspaper articles over a comparable time period (1^st^ Jan 2001-31^st^ May 2021 inclusive). Full details of the methodology are outlined in File S1. A list of the details of resulting newspaper articles will be available to download from the LJMU data storage repository.

To provide a comparison with peer-reviewed publications, we examined newspaper articles applying the same five framing categories to the entire body of text. Additionally, we applied inductive processes to classify further themes that differed from those observed within the peer-reviewed literature, resulting in five additional inductive categories: ‘Laws and Regulations’, ‘Public Health and Safety’, ‘Irresponsible Pet Ownership’, ‘Illegal Trade’, and ‘Defence of Trade’. We acknowledge that these choices (and the framing categories used for peer-reviewed publications) may be informed by our own experiences, knowledge, and culture, and could vary according to the people developing them. We are confident, though, that in our categorisations we have captured the main themes related to the exotic pet trade given the existing biases in the literature. Frequency of newspaper articles within a framing category (‘Files’), and frequency of mentions within the article (‘References’) were calculated for each frame, and the content of each article extracted for further analysis regarding language use and narrative. Thematic analysis comprised ‘coding’ themes within newspaper articles, meaning that relevant words were not extracted in isolation, but sections of text were coded to provide context. The approach to coding and analysis was partially grounded, as we were influenced by knowledge and personal experience; however, the inductive coding reflected framing not found in the deductive coding, highlighting that these patterns were mostly absent from peer-reviewed publications. The codebook was developed by SW upon reading all articles to gain a sense of their content and tone and defining themes. SW and GEA coded the text according to these themes, clarifying any ambiguities in code descriptions and highlighting any potential themes that had been missed. This iterative process continued until the authors were satisfied that the codebook reflected the themes relevant to our questions. The final codebook was then used to recode the same set of articles, and a random sample of articles (∼10%) were cross-checked between these two authors.

### Comparisons and statistical analyses

#### Frequency of frames and taxonomic foci in peer-reviewed literature

G-tests were used to analyse variation in the number of peer-reviewed publications in each taxon:framing combination. Distributions of the count of peer-reviewed publications per year were non-normal. Consequently, to analyse whether they varied with each of framing category and taxonomic focus separately we used non-parametric Kruskal-Wallis tests followed by Dunn’s multiple comparison *post hoc* tests. All statistical analyses were conducted in the software package R (R Core Team 2022).

#### Factors associated with citations per year

We calculated the number of citations per year to be used as a response variable in a mixed-effects model. Year of publication was included as a random effect, and framing category and taxonomic focus were included as fixed effects. We also included as fixed effects the impact factor of the journal, and the h-index of the lead author as they are likely to play an important role in the citation rate. Mean citations per year and journal impact factor were log transformed to make the distribution of the residuals meet the assumptions of the model.

Model outputs were compared using Akaike information criterion (AIC) and models within 6 units of each other were considered to be equally supported (Richards 2005). Individual variable importance judged based on the parameter estimate effect sizes and t-values in the model.

### Comparison of language between the two media-types

The tidytext package provides access to two general-purpose lexicons: afinn (Bradley & Lang 2009), and nrc (syuzhet; Mohammad & Turney, 2012). Both contain many English words and are based on single words, which are assigned scores for positive/negative sentiment. The afinn lexicon assigns words with a score of -5 to 5, with negative scores indicating negative sentiment and positive scores indicating positive sentiment. The nrc lexicon categorizes words into the following: positive, negative, anger, anticipation, disgust, fear, joy, sadness, surprise, and trust. Both lexicons were implemented in this project, to maximise opportunity to explore variation in measures of sentiment between framings and media types. For newspaper articles, the whole body of text was analysed in this way, whereas for the peer-reviewed publications only the abstract was used. Packages tidytext (Silge & Robinson 2016) and tm (Feinerer et al. 2008) were used for text mining. Figures were produced using ggplot2 (Wickham 2016).

## Results

Our search resulted in 320 suitable peer-reviewed publications focused on the exotic pet trade between 2001 and 2021 (Fig S1). Data from 2021 were excluded from any analyses relying on a full year of data (e.g., papers per year and citations per year), resulting in n=305 for those. Peer-reviewed publications per year showed a large increase in number ranging from single figures in 2011, to a high of 53 in 2020 (Fig 1), and they were not distributed evenly across framings. Across the entire study period the following frequency of framings were observed: 55.3% ‘Multiple frames’ (n=177), 19.1% ‘Conservation’ (n=61), 7.8% ‘Disease’ (n=25), 7.5% ‘Invasive species’ (n=24), 6.6%, ‘No framing’ (n=21), 3.1% ‘Welfare’ (n=10), 0.6% ‘Economics’ (n=2). Within the 177 ‘Multiple frames’ peer-reviewed publications eleven combinations were present: with the most frequent combinations including Conservation-Invasion (n=29), Conservation-Economy (n=21), and Disease-Welfare (n=10). In addition, 45.2% (n=80) of the 177 ‘Multiple frame’ papers included three or more framings. A full list of all combinations of framings and their frequencies are available in Table S5.

**Figure 1:**
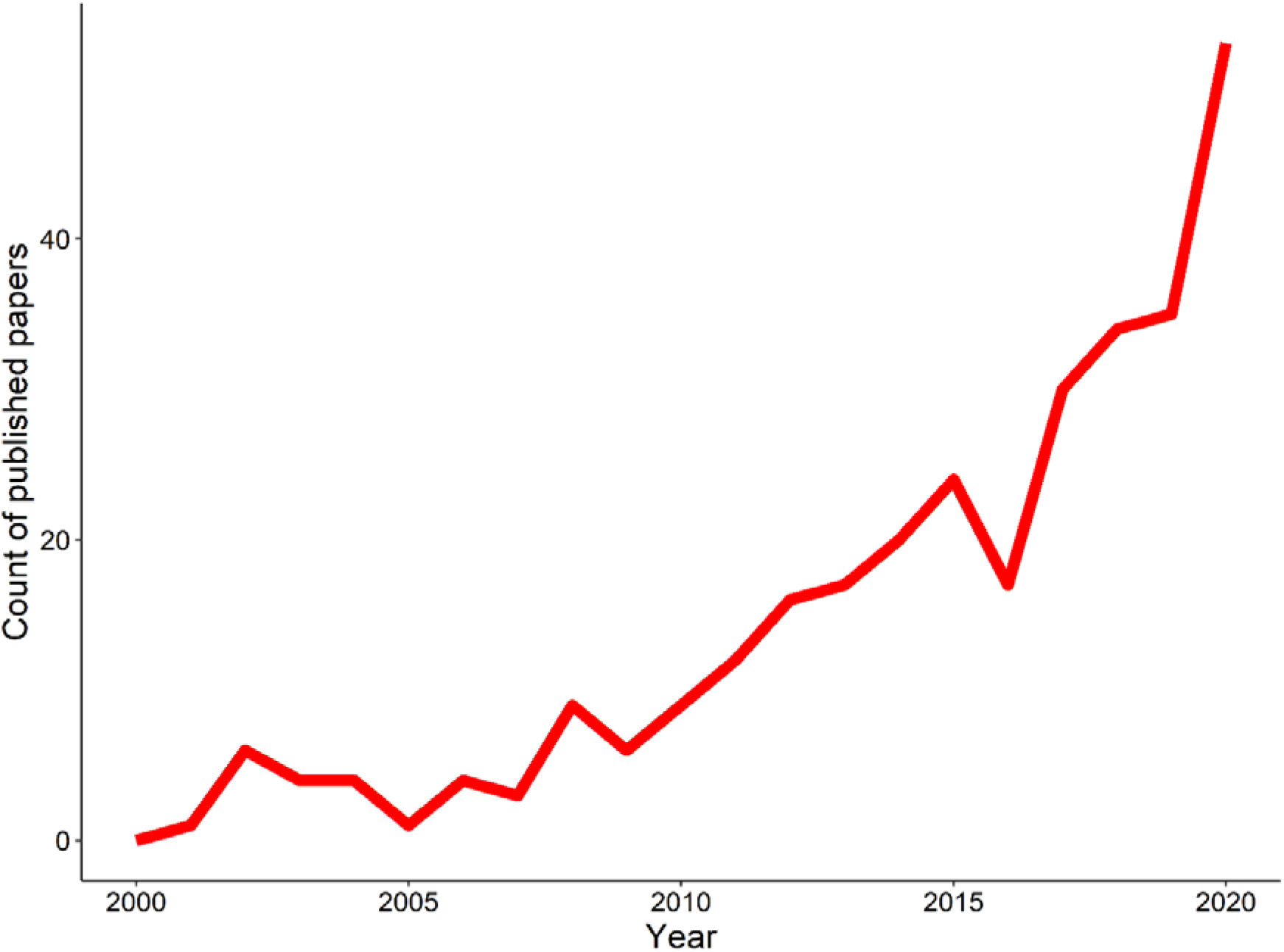
Number of peer-reviewed papers published per year, focusing on the exotic pet trade, between the years 2000 and 2020 (n=305).

Peer-reviewed publications were not evenly distributed across combinations of framing and taxon (Table 1), both including data on a single category/focus (G=36.89, Χ^2^ df=16, p=0.002) and those within multiple categories/foci (G=78.97, Χ^2^ df=36, p<0.001). The largest general patterns to be noted are: (1) most peer-reviewed publications were framed in multiple ways, featuring multiple taxonomic groups, and (2) relatively few were framed within a singular category focussed on ‘Welfare’ or ‘Economics’.

**Table 1:**
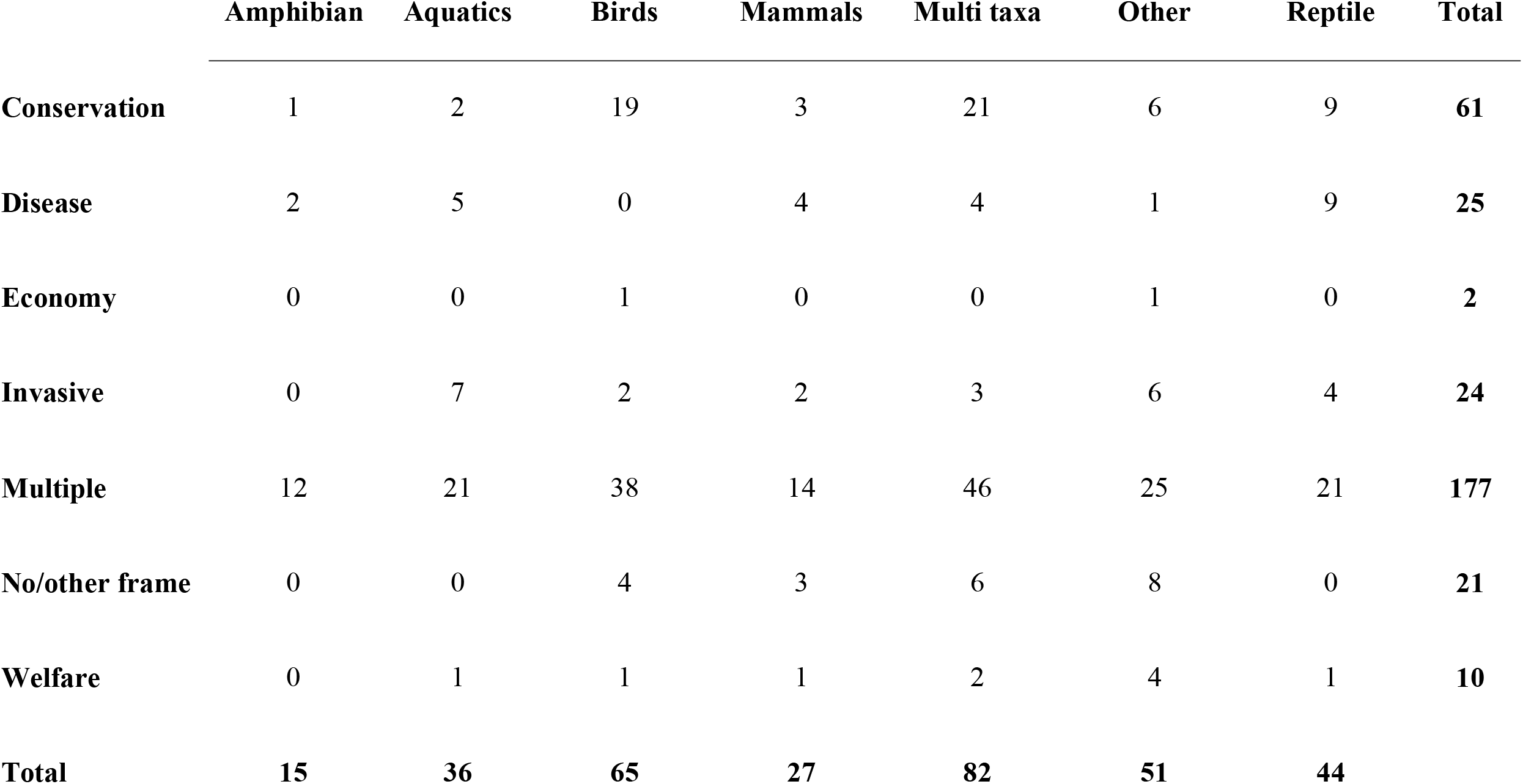
Raw count of peer-reviewed publications in each combination of the framing categories and taxonomic foci.

|  | **Amphibian** | **Aquatics** | **Birds** | **Mammals** | **Multi taxa** | **Other** | **Reptile** | **Total** |
| --- | --- | --- | --- | --- | --- | --- | --- | --- |
| **Conservation** | 1 | 2 | 19 | 3 | 21 | 6 | 9 | **61** |
| **Disease** | 2 | 5 | 0 | 4 | 4 | 1 | 9 | **25** |
| **Economy** | 0 | 0 | 1 | 0 | 0 | 1 | 0 | **2** |
| **Invasive** | 0 | 7 | 2 | 2 | 3 | 6 | 4 | **24** |
| **Multiple** | 12 | 21 | 38 | 14 | 46 | 25 | 21 | **177** |
| **No/other frame** | 0 | 0 | 4 | 3 | 6 | 8 | 0 | **21** |
| **Welfare** | 0 | 1 | 1 | 1 | 2 | 4 | 1 | **10** |
| **Total** | **15** | **36** | **65** | **27** | **82** | **51** | **44** |  |

The number of peer-reviewed publications per year differed significantly with framing (Χ^2^=47.22, d.f.=6, p<0.001; Fig 2). The most common framing was ‘Multiple frames’ (median=6.00, IQR=9.50), which was used significantly more than all other framings, aside from those framed in the context of ‘Conservation’ (median=3.00, IQR=2.50). Those framed solely around ‘Economics’ (median=1.00, IQR=0.25) and ‘Welfare’ (median=1.00, IQR=0.00) were significantly less common than those based on ‘Multiple frames’ or ‘Conservation’ alone.

**Figure 2:**
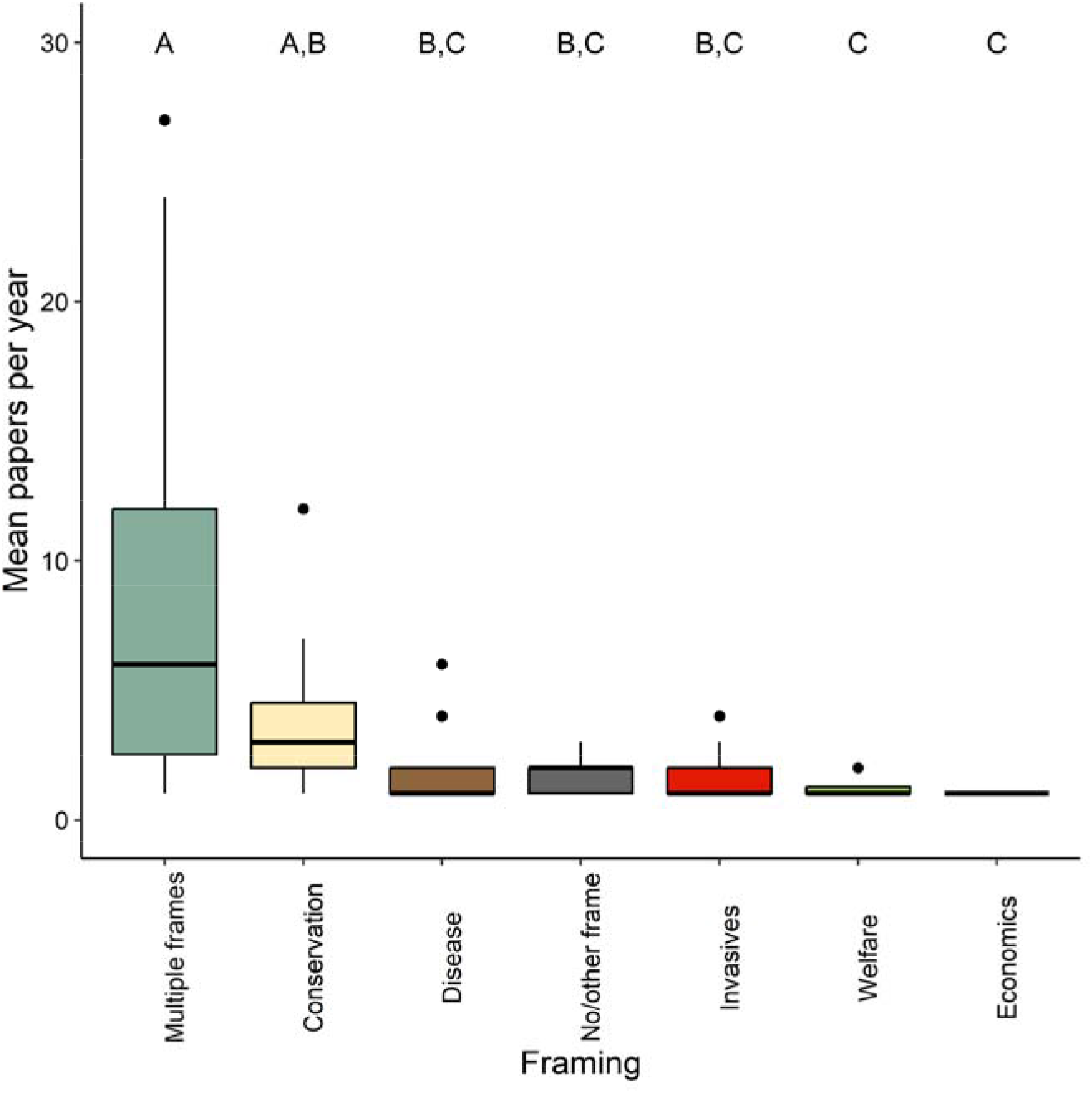
Mean peer-reviewed papers, per year, including the framing categories: Conservation, Disease, Economics, Invasives, Welfare, Multiple frames, or No/Other frame. Upper case letters indicate the grouping of framings based on significant differences found in a Kruskal-Wallis test.

The number of peer-reviewed publications per year differed significantly among taxa (Χ^2^=14.56, d.f.=6, p=0.024; Fig 3). They were more frequently focussed on ‘Multiple taxa’ (median=4, IQR=3.5) or ‘Birds’ (median=4, IQR=6), than solely on ‘Aquatics’ (median=1, IQR=2), ‘Mammals’ (median=1, IQR=2), or ‘Amphibians’ (median=1, IQR=0.5). Those that focused on ‘Amphibians’ were also less common than peer-reviewed publications with no taxonomic focus (median=2.5, IQR=2). There were no other significant differences amongst categories.

**Figure 3:**
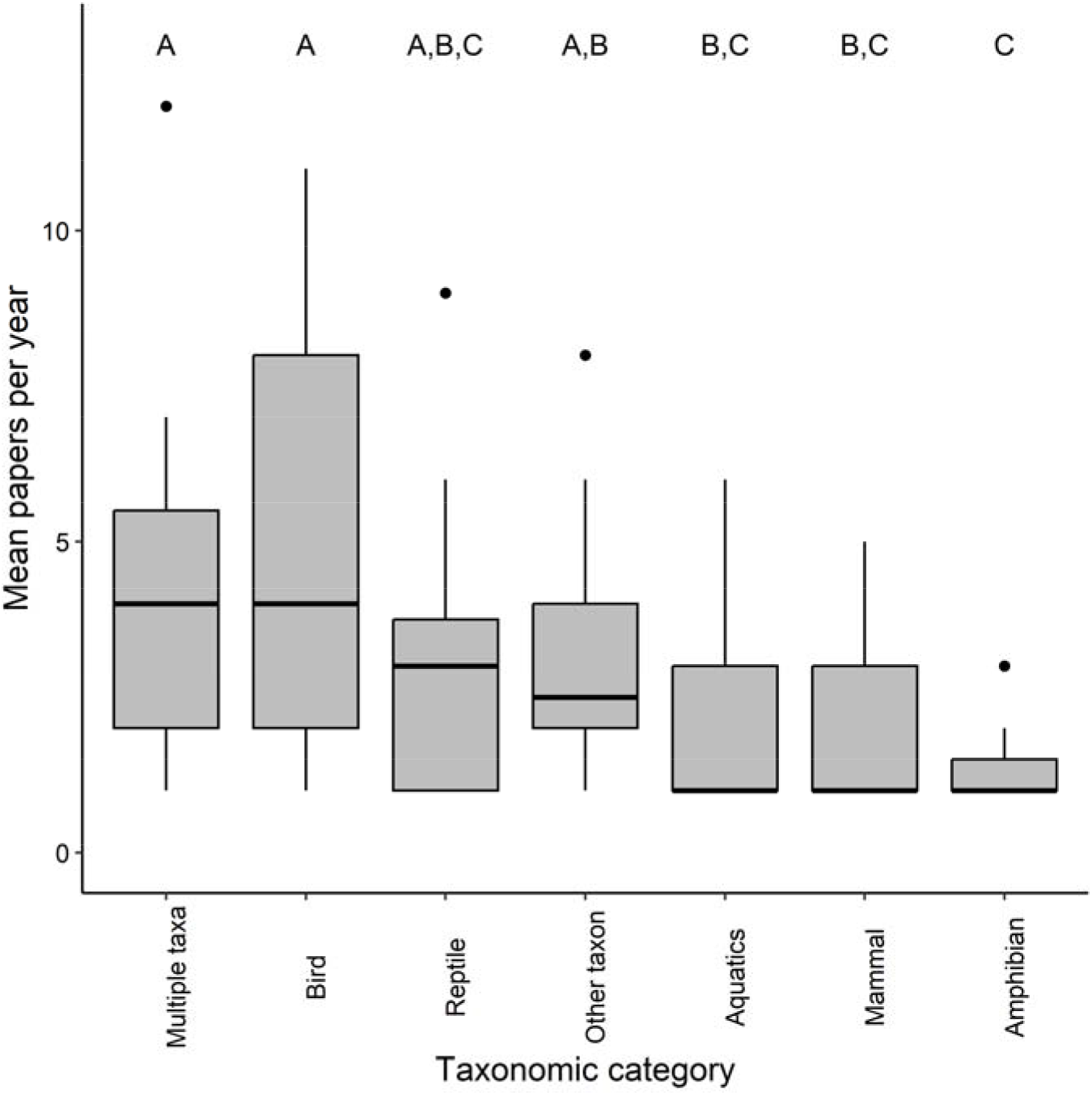
Mean peer-reviewed papers, per year, in each of the five taxonomic categories: Amphib (i.e., Amphibians), Aquatics, Birds, Mammals, and Reptiles – and Multiple taxa or Other taxon. Upper case letters indicate the grouping of framings based on significant differences found in a Kruskal-Wallis test.

The top performing model of citations per year contained the h-index of the first author, and the impact factor of the journal (AIC of 801.72; Table 2). The next best performing model had a ΔAIC of -4.46 relative to the best model, which suggests an equal level of support for both. This model additionally contained a term for the framing category suggesting an important role for this variable in determining the citation rate. The next best performing model was almost 9 AIC units higher (8.93), suggesting a large drop-off in model support.

**Table 2:** Linear mixed model outputs showing relative performance of models explaining the number of citations per year an article received (within peer-reviewed scientific literature with ‘year’ as a random effect).

| Model | d.f. | AIC | ΔAIC | R<sup>2</sup> |
| --- | --- | --- | --- | --- |
| **First Author h + Impact Factor** | **5** | **801.72** | **-** | **0.34** |
| **Framing + First Author h + Impact Factor** | **11** | **806.18** | **4.46** | **0.36** |
| Framing + Taxon + Impact Factor | 16 | 815.11 | 13.39 | 0.36 |
| Taxon + First Author h + Impact Factor | 11 | 818.56 | 16.84 | 0.34 |
| Framing + Taxon + First Author h + Impact Factor + Framing*Taxon | 41 | 821.15 | 19.43 | 0.37 |
| Framing + Taxon + First Author h + Impact Factor | 17 | 822.94 | 21.22 | 0.37 |
| Framing + Taxon + First Author h + Impact Factor + Impact Factor*Framing | 23 | 831.00 | 29.28 | 0.37 |
| Framing + Taxon + First Author h + Impact Factor + Framing*First Author h | 23 | 853.43 | 51.71 | 0.38 |
| Framing + Taxon + First Author h | 16 | 876.83 | 74.81 | 0.18 |

Parameters of the best performing model suggest that peer-reviewed publications framed around welfare are cited fewer times per year than some other framings (Table 3, Fig 4). For example, ‘Multiple frame’ peer-reviewed publications were cited more frequently that other categories, with a four-fold difference between them (mean=3.44 citations per year) and the least cited framing (‘Welfare’, mean=0.75 citations per year).

**Figure 4:**
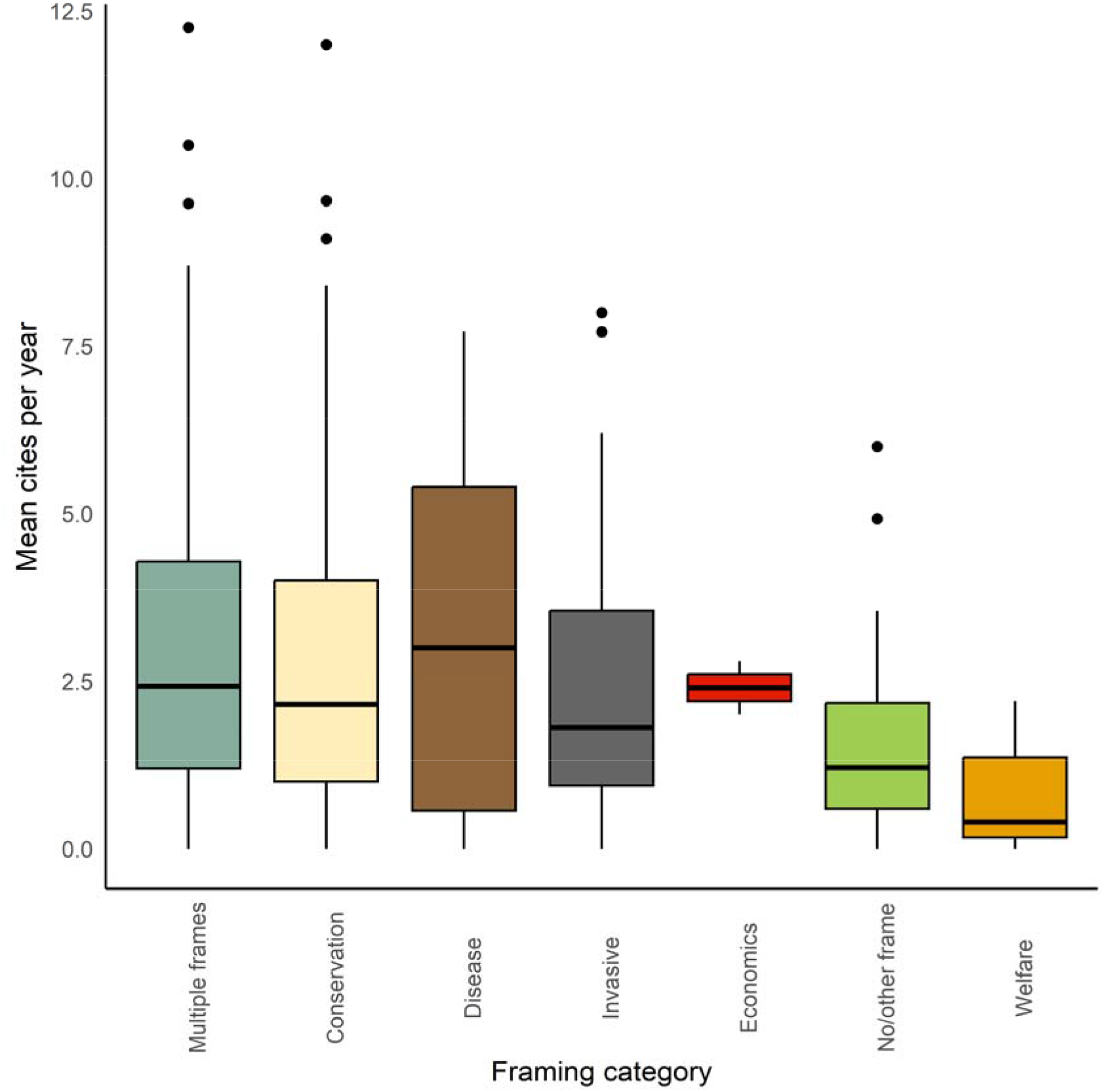
Mean citations for peer-reviewed papers, per year, for each of the five framing categories: Conservation, Disease, Invasives, Welfare, and Economics - and those categories Multiple frames and No/other frame.

**Table 3:**
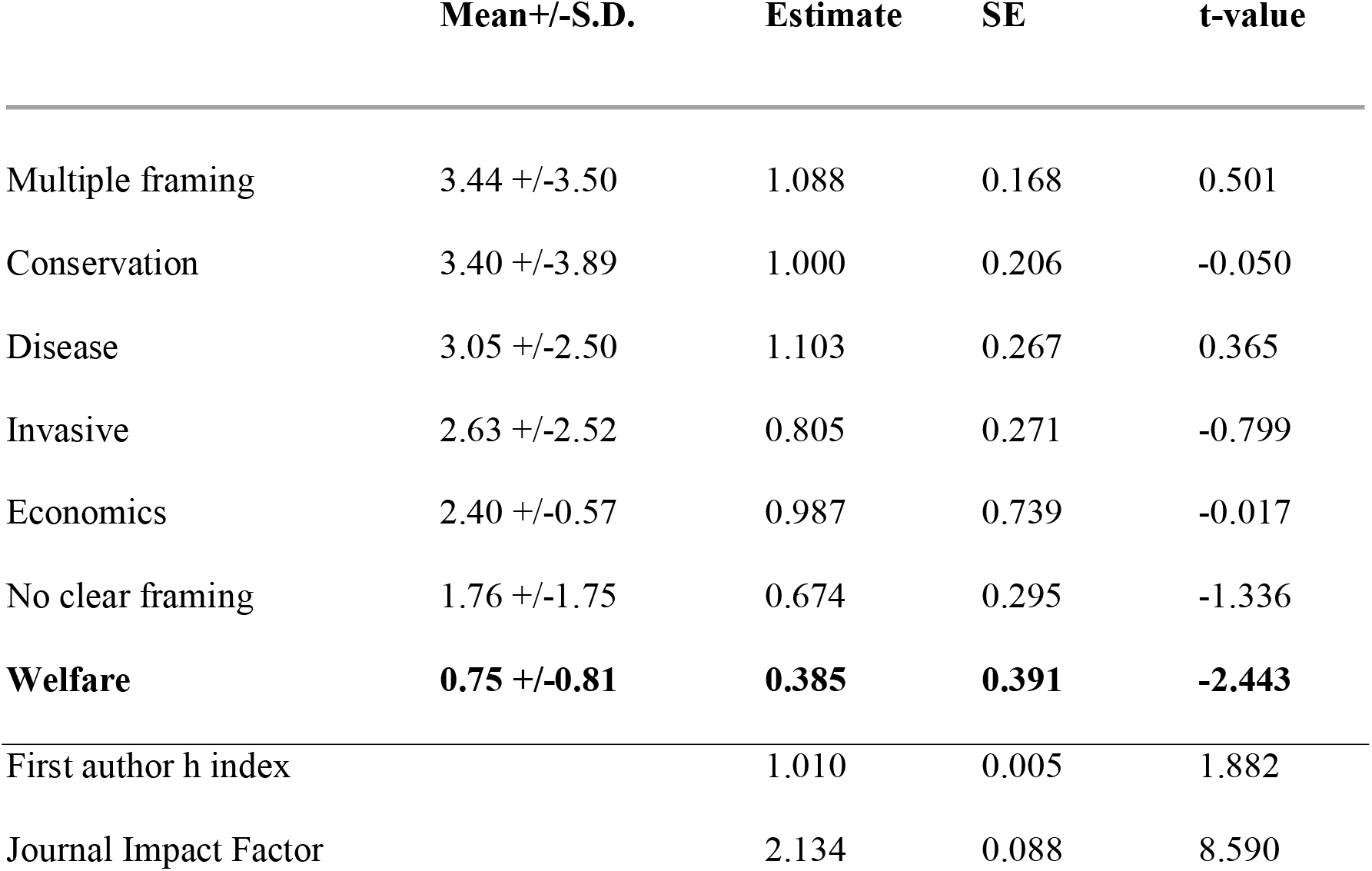
Mean and S.D. values of peer-reviewed paper citations per year for each framing category, along with the parameter estimate for each from the best performing model. Framing categories are ordered in descending order of mean citations per year and those with t-values exceeding 2 are in bold.

191 newspaper articles were included in our analyses. Of the five inductive themes used to categorise them ‘Welfare’ was the most common framing and had the most references to that framing within a given article (Table 4). The frequency with which ‘Welfare’ and ‘Economy’ were used as a framing within newspaper articles compared to peer-reviewed publications were the most marked difference between the two media-types (Fig 5).

**Figure 5:**
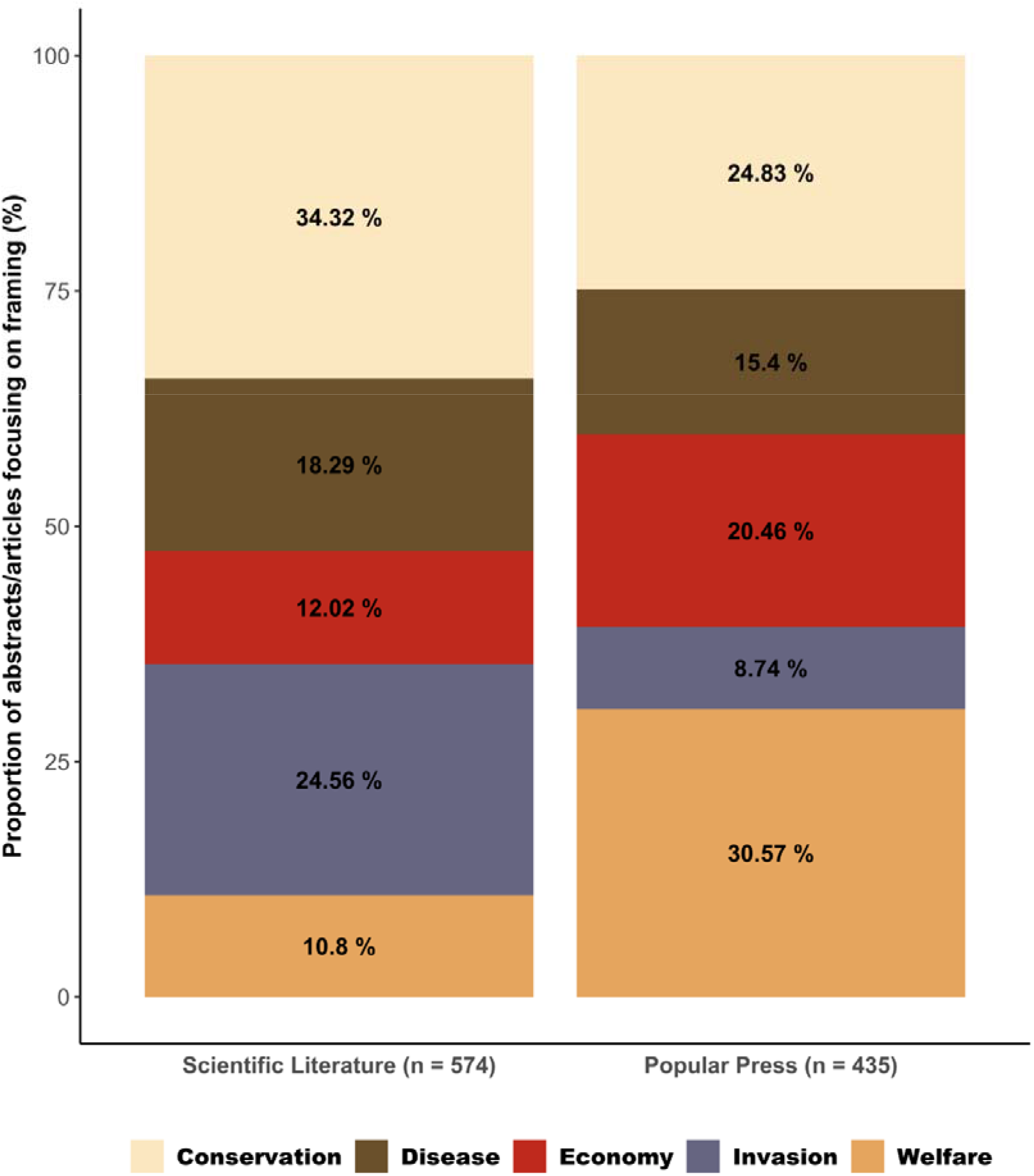
Proportion of each of the five framings in peer-reviewed scientific literature abstracts and newspaper articles. Abstracts/articles included in more than one framing are included in all for the purpose of this data visualisation.

**Table 4:**
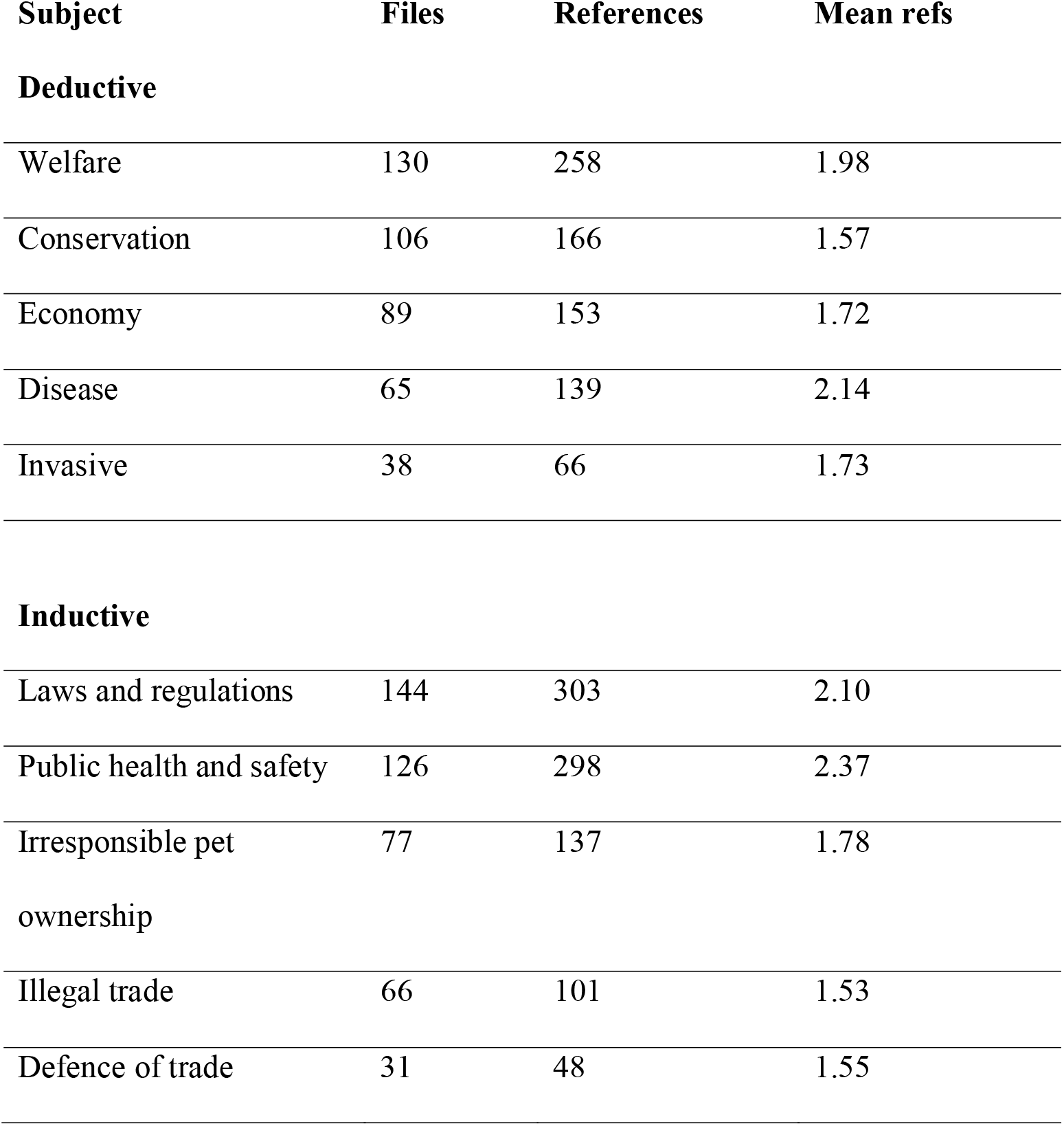
Quantitative analysis of deductive and inductive coding from newspaper articles (*n*=191). ‘Files’ refers to the number of articles in which the theme occurred, ‘References’ are the total number of items coded to that theme. A single news article may contain more than one of the subjects listed below.

‘Conservation’ was proportionally the highest framing in peer-reviewed publications (34.3% of abstracts) and second highest in newspaper articles (24.8% of articles), but still suffered a percentage point reduction of almost 10% between the two media-types. In contrast, ‘Welfare’ was the least frequently used frame in peer-reviewed publications and highest in newspapers (10.8% compared to 30.6%), and represents the largest change of any frame between the two media-types. When deductive categories were also included, the most frequent framings in popular media were ‘Laws and Regulations’ and ‘Public Health and Safety’, both of which were used as framings at a comparable frequency to ‘Welfare’ in the inductive categories. For newspaper articles within these framings the topics were also referenced frequently (Table 4).

Afinn sentiment analyses (Fig 6) suggests that the language used in newspapers was substantially more negative than that used in the abstracts of peer-reviewed papers. Within a single media-type, there were also notable differences in language use between framing categories. For example, within abstracts framed around ‘Welfare’, there was less negative language than other categories, particularly those framed around ‘Conservation’, ‘Disease’, and ‘Invasive species’. In newspapers articles the differences in sentiment between categories was relatively smaller.

**Figure 6:**
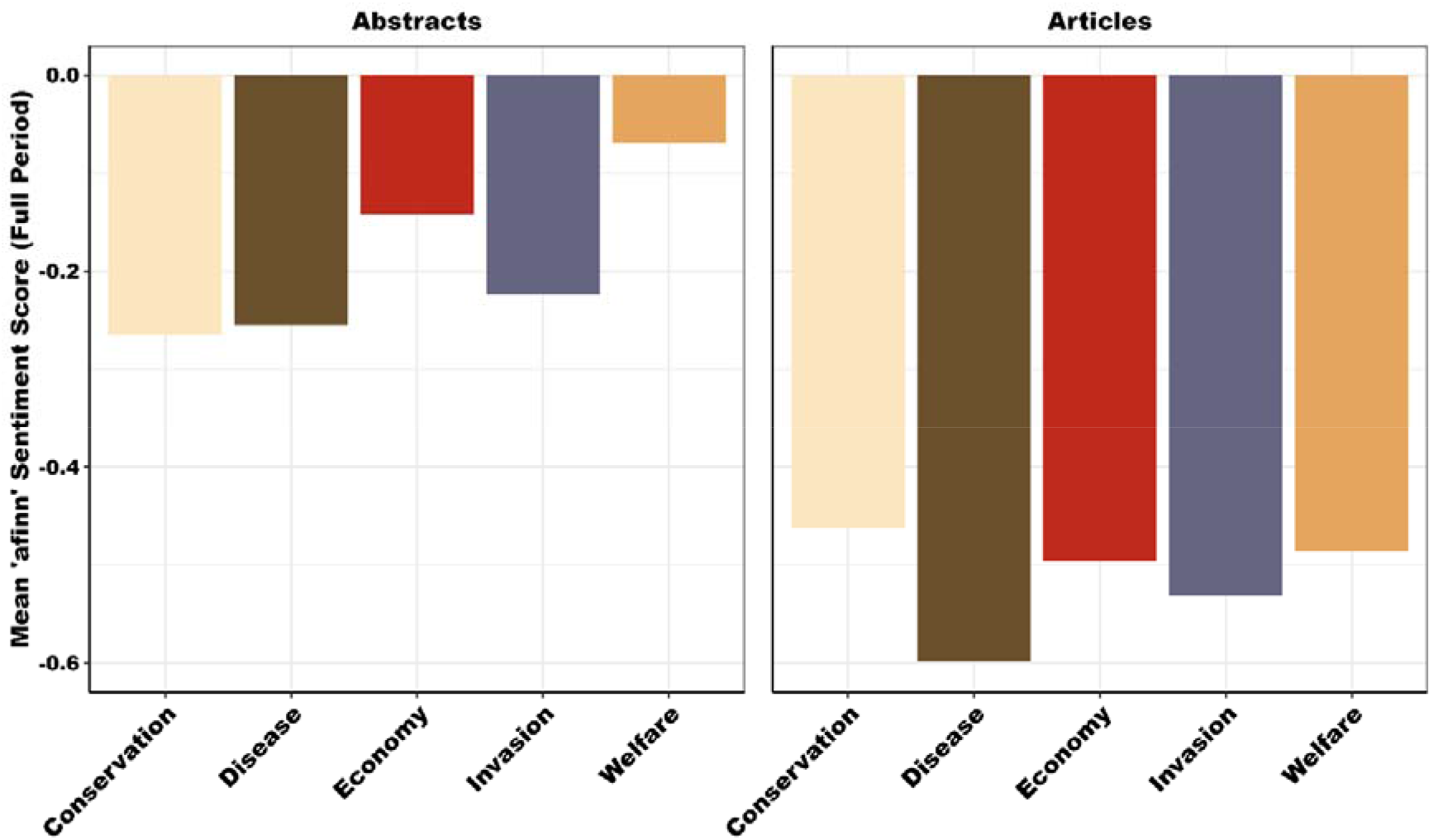
Mean afinn sentiment scores among the five framing categories for peer-reviewed abstracts (top) and newspaper articles (bottom). Peer-reviewed abstracts typically contained less negative language from the afinn lexicography than newspaper articles. There was relatively more variation in sentiment amongst framings in peer-reviewed abstracts, with economics and welfare being less negative than other framings.

Nrc sentiment scores suggest some consistency in emotive language category use for a given framing, and among framing types within a media-type. However, at the broadest scale between media-types there is a great deal of variation in language category used. For example, within newspaper articles the following language types are greater than that seen in abstracts: ‘Anger’, ‘Sadness’, ‘Fear’, ‘Joy’, and ‘Anticipation’ (Table 5). Peer-reviewed abstracts are higher for ‘Trust’ and ‘Positive’ language. There is not variation between the two media types for ‘Negative’ language, ‘Disgust’ or ‘Surprise’. Examples of a selection of articles scoring highly for these categories are available in S6 Table.

**Table 5:** Comparison of percentage of newspaper articles and peer-review publication abstracts consisting of each of the nrc language type categories. We have defined there to be a difference between the two media types when the range of percentages do not overlap between the media types.

|  | Articles (%) | Abstract (%) | Variation? |
| --- | --- | --- | --- |
| **Anger** | 6.2-6.5 | 4.7-5.3 | Newspaper articles higher % |
| **Anticipation** | 8.2-8.5 | 7.0-8.1 | Newspaper articles higher % |
| **Fear** | 11.4-12.1 | 8.1-11.0 | Newspaper articles higher % |
| **Joy** | 5.2-5.8 | 3.9-4.7 | Newspaper articles higher % |
| **Sadness** | 7.1-7.6 | 5.0-6.9 | Newspaper articles higher % |
| **Positive** | 20.4-21.2 | 24.2-27.6 | Peer-review abstracts higher % |
| **Trust** | 12.0-12.6 | 14.5-17.5 | Peer-review abstracts higher % |
| **Disgust** | 4.7-5.7 | 3.7-5.0 | Overlaps – no difference |
| **Negative** | 18.2-19.0 | 16.9-18.6 | Overlaps – no difference |
| **Surprise** | 3.7-4.0 | 3.8-4.1 | Overlaps – no difference |

The shape of the profiles of relative use of the categories are largely consistent (Fig 7). In all 10 combinations of framing and media-type, the five most used categories of words were consistent. In all 10 combinations of framing and media-type, the three most used language categories were used in over 50% of the words in those articles.

**Figure 7:**
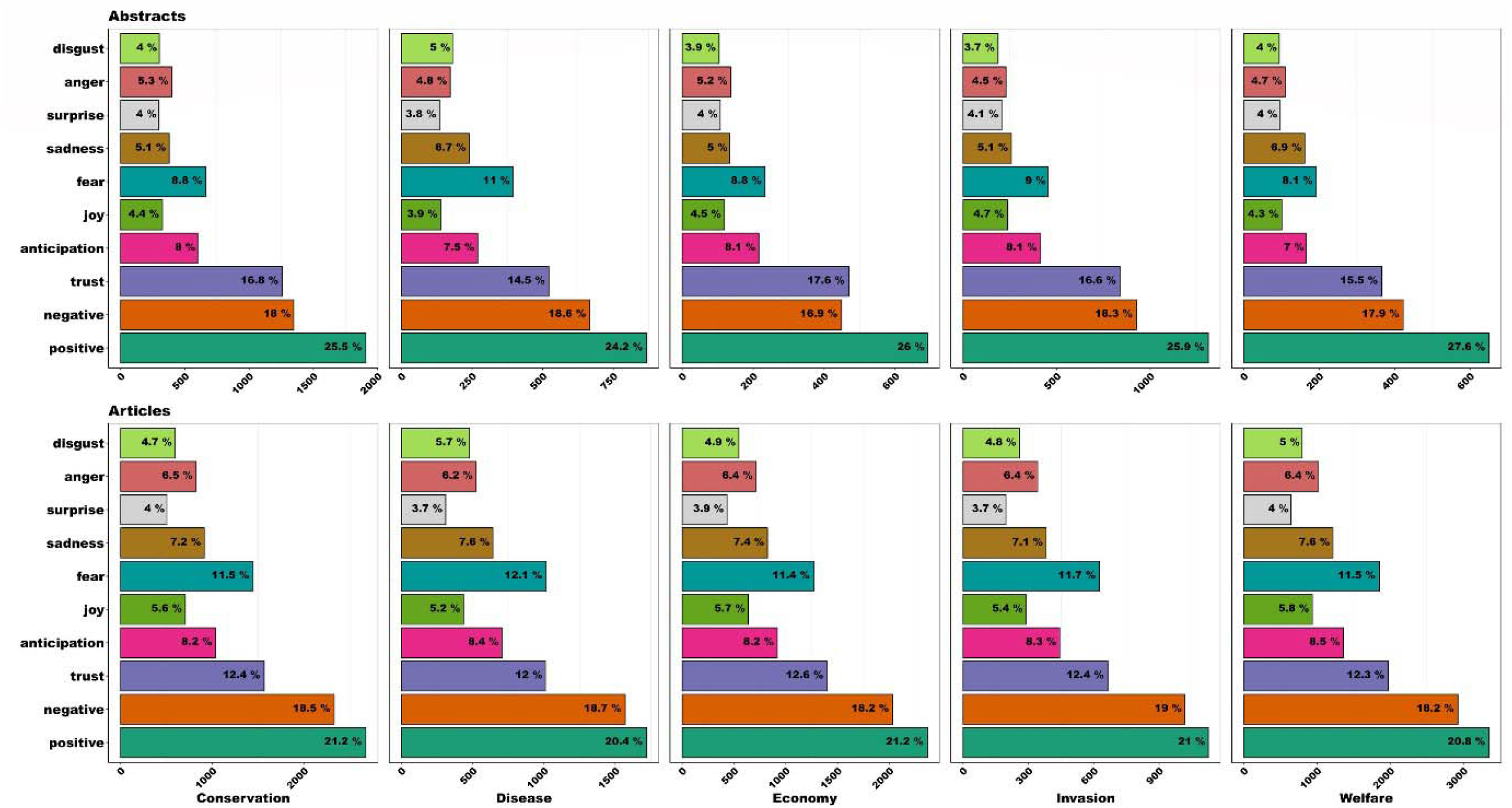
nrc sentiment scores (%) for all five framings for both peer-reviewed abstracts (top) and newspaper articles (bottom). Profiles and relative ranks of each of the eight emotions and two sentiments (positive/negative) encoded in the nrc lexicography are consistent across all combinations of framing and media-type. Note the x-axis changes scale as the absolute number of words varies between framings.

## Discussion

The number of peer-reviewed publications focusing on the exotic pet trade has increased greatly since the turn of the Century. Our analyses suggest that peer-reviewed papers on the exotic pet trade were frequently framed around multiple topics and focussed on multiple taxa. In single-frame peer-reviewed papers there was a clear contrast with newspaper articles in that welfare-based articles were less frequent and less well-cited, whereas newspaper articles were framed around welfare more commonly than other frames. There were also differences in language used between framings and media-types, although the precise differences were context dependent, varying with the metric of language sentiment implemented. The potential drivers of these patterns, and their implications are discussed below.

Peer-reviewed papers with multiple framing foci were not only more frequent than other categories but also had a higher rate of citations per year. Given the complexities of modern global society, the need for a more systems-based approach is being recognised in strategies for conservation (One Plan; CBSG 2023), ecosystem health (OneHealth; WHO 2023), metrics of economic health (Doughnut Economics 2023), tackling wildlife trafficking (Gore et al. 2023), and living collections (Spooner et al. 2023). As a large proportion of papers in the peer-reviewed literature were framed in this multi-faceted manner and focussed on multiple taxa, it may be hoped that a broad, inclusive evidence-base for the effects of the exotic pet trade is being developed. Unfortunately, recent research suggests that significant geographic, taxonomic, and economic knowledge gaps exist in our understanding of the trade, leading to calls for more coordinated, structured approaches (Sinclair et al. 2021; Keskin et al. 2023; Sengottuvel et al. 2023) that may incorporate multidisciplinary teams to optimise their chance of success (Gore et al. 2023).

The current biodiversity crisis may explain why conservation-based peer-reviewed research was both highly prevalent and highly cited. However, in many cases, despite the scale of trade, there remains a lack of empirical data linking traded animals to population viability (see Marshall et al. 2020. Natusch et al. 2021a and b; Sosnowski and Petrossian 2021), which is a stumbling block to those advocating sustainable use of animals for the pet trade (Natusch et al. 2019). Under sustainable-use models, components of biodiversity are used at a rate that does not lead to its long-term decline, and therefore maintains its potential to meet the needs of present and future generations (IPBES 2019). The strategy can be particularly important for supporting local livelihoods and engendering equity of roles, rights, and agency of peoples within range states (Obura et al. 2023). However, there are relatively few empirical examples linking the pet trade to the benefit of member state communities or written within the context of longer-term sustainability of traded populations (CITES and Livelihoods 2023, Species Use Database 2023). Where such studies do exist, they often mention the limited scope of the pet trade in poverty alleviation, or an associated lack of motivation for effective stewardship of traded species (Robinson et al. 2018a and 2018b). The latter could be reflected in our data by the small number of peer-reviewed papers framed within economics and the relatively low citation rate of those that do exist. Given the importance of the global pet trade to diverse stakeholder groups throughout the supply chain (Sinclair et al. 2021), the lack of peer-reviewed literature on the sustainability and economic benefits of the trade highlights a key knowledge gap.

Establishing a stronger evidence base on community-led sustainable-use initiatives could better balance the cultural and socioeconomic benefits of the pet trade against the risks it represents. However, at present the dominance of English as the language of scientific discourse, evaluation, and knowledge sharing could provide a major stumbling block to such evidence reaching the peer-reviewed literature (Khelifa, et al. 2022; Amano et al. 2023). In this study we focussed on English language articles because of their historic and present dominance in the peer-reviewed field, but this has resulted in significant biases in a researcher’s ability to access and/or disseminate their scientific knowledge and costs to the researchers involved (Ramírez-Castañeda 2020; Amano et al. 2023). Constructive solutions for making the field more equitable, tapping into non-English science, and solving this language barrier could provide long-term solutions to some of these inequities (Amano et al. 2021; Khelifa, et al. 2022). Analyses such as ours conducted at a finer-scale and in native language media within key sink and source regions would likely highlight context-dependent results and potentially identify important regional variation in media coverage of the subject matter. These analyses could highlight cultural similarities (Haddon et al. 2021, De la Fuente et al. 2023) and differences in attitudes towards keeping exotic pets, fundamental concepts related to the trade, and how they may change over time (Guo and Meijboom 2023, Mata, et al. 2023a; Mata, et al. 2023b). Any such cultural and demographic differences must be acknowledged and understood if multination systems-based approaches to managing the global pet trade are to be successful.

The lack of focus on welfare in peer-reviewed papers is consistent with previous research (Baker et al. 2013) suggesting that relatively little progress has occurred in this area. As our understanding of animal cognition and sentience across the animal kingdom developments and as associated legislation changes accordingly (Browning and Veit 2022), it is imperative that to optimise their welfare, we incorporate this information into practices that use live animals. The results of our analyses suggest that, as of 2021, very little progress has been made in linking the exotic pet trade with our improving knowledge of animal welfare. Measuring and incorporating the needs and requirements of traded animals at each stage of the supply chain, from their origin to how they are kept as pets, is a key challenge for peer-reviewed science, and one that we appear to be failing to meet. In contrast, while peer-reviewed publications exhibited a severe shortfall in welfare-based studies, newspaper articles were most frequently framed in the context of animal welfare.

So why would welfare vary so greatly in its coverage by these media types, and why does language use between them differ so much? Some of this may be due to the structure and nature of the media in question. Newspaper articles can take many different structures, but will often include a ‘story’ element, which is designed to elicit an emotional response (Dahlstrom 2014). Stories will commonly focus on an individual (whether it be a human or an animal; Wallack and DeJong 1995), will have a beginning and middle and an end, and will often be aligned to a specific objective or brand (Galtung, Ruge 1965). In contrast, peer-reviewed articles are typically more neutral in nature and should, in theory, be focussed on the internal value of the information within, rather than appealing to an emotional response from the readers and will often specifically avoid the central characteristics of storytelling (Katz 2013; examples of newspaper article and peer-reviewed abstract on a similar subject are available in the File S1 section “Example of narrative-driven newspaper article versus data-driven peer-reviewed abstract.”). In the case of animal welfare, it is relatively easy to understand how an emotive narrative-based piece could be written around the experience of a pet or its guardian. Welfare may therefore be perceived to be an individual-level phenomenon. In contrast, conservation, in which attention and action are usually focussed on the level of the species or population (Shaffer 1981), is, for that very reason often seen to be a broader scale issue. The tendency for newspaper articles to be based around individual-level interests may also explain why analysis of newspaper articles identified commonly used frames that were not found in peer-reviewed papers: ‘Laws and Regulations’ and ‘Public Health and Safety’. For example, emphasis on information related to the risk of infectious disease and legality of the pet trade has been linked to a reduction in the likelihood of consumers purchasing exotic pets (Moorhouse et al. 2017), highlighting the individual-level influence these framings can have, and hence their popularity as a framing tool in newspaper articles.

The mismatch between media types in coverage of welfare could also be indicative of a significant difference in what funding bodies (and therefore peer-reviewed journals) view as priority research directions, and those that newspaper articles indicate as being important to the public. While research funding priorities should not be the subject of a popular vote, funding priorities and policies vary greatly and can be extremely complex in terms of who they are aligned with and driven by (Stimson et al. 1995, Jacobs and Page 2005, Steinberg 2011). If we do want to develop more systems-based approaches to resolve issues related to the exotic pet-trade, more strategic, multidisciplinary funding priorities to do so would be a step forward in this process.

Selectivity in the framing of a topic communicated via any channel can cause inconsistencies between the prominence of the subject matter and its relative importance as an issue. Recent research has highlighted that there are mismatches between media coverage, public engagement, and scientific consensus in several environmental contexts (Hammond et al. 2022, Santos and Chowder 2021, Shiffman et al. 2020, Walker et al. 2019). These mismatches can be important as both scientific evidence and public engagement are essential when developing policy and legislation. A future challenge for both media avenues is to communicate clearly our current state of knowledge and where gaps lie, whilst continuing to engage readers with longer-term societal issues (e.g., climate change and the biodiversity crisis). With a view to making peer-reviewed science more inclusive, the scientific community is currently undergoing changes in the way it works by taking steps to make scientific knowledge open access and therefore more inclusive (Ross-Hellauer et al. 2022). Another step along this road to inclusivity would be to better bridge the gap between the people generating that knowledge and those who either use it, or those who may be affected by decisions informed by it. Part of the solution could be better use of knowledge brokers (Meyer 2010), who are trained to move information between researchers and practitioners, policymakers, and those affected by policy. Another improvement might to be consider the intellectual accessibility of science in terms of how language-use and structure facilitates or hinders understanding and engagement with a broader audience (Dahlstrom 2014). Combined, these changes could provide decision makers and voters with the means to marry empirical evidence with ethical standpoints and public engagement when shaping policy and legislation within the exotic pet trade.

## Author Contributions

All authors conceived the ideas and designed methodology; SW collected the data; All authors analysed the data; JB and SW led the writing of the manuscript. All authors contributed critically to the drafts and gave final approval for publication.

## Acknowledgements

This study was partly funded by a Liverpool John Moores Impact Grant. The authors have no conflicts of interest to declare.

## Supporting Information

Table S1: Details of terms used for peer-reviewed scientific literature search. Note: words containing * allow for variants to appear in search results, including pluralisation.

Table S2: Details of ‘pet trade’ and related terms for secondary screening of articles to ensure a focus on the exotic pet trade.

Table S3: Details of each framing category and related terms.

Table S4: Details of taxonomic foci and related terms.

Table S5: Raw count of peer-reviewed papers in each framing category and combinations thereof.

Table S6: Examples of a selection of articles scoring highly for different categories of NRC emotion categories.

Fig S1: PRISMA flowchart showing the number of peer-reviewed articles at each point in the screening process.

File S1: Supplementary information on peer-reviewed article screening process and examples of articles with different language use and narration style.

## Notes

### Competing Interest Statement

The authors have declared no competing interest.

## References

Alves RRN, et al. 2012. A review on human attitudes towards reptiles in Brazil. Environmental monitoring and assessment. 184, pp.6877–6901. 10.1007/s10661-011-2465-0 PMID: 22134858

Alves RRN. 2012. Relationships between fauna and people and the role of ethnozoology in animal conservation. Ethnobiol Conserv.1:12–69.

Alves RRN, Rocha LA. Fauna at home: Animals as pets. In Ethnozoology (pp. 303–321). 2018. Academic Press.

Amano T, et al. 2021. Tapping into non-English-language science for the conservation of global biodiversity. PLoS Biology 19(10) p.e3001296.

Amano T, et al. 2023. The manifold costs of being a non-native English speaker in science. PLoS Biology 21(7) p.e3002184.

Baker SE, Cain R, Van Kesteren F, Zommers ZA, D’Cruze N, Macdonald DW. 2013. Rough trade: animal welfare in the global wildlife trade. BioScience 63(12) 928–938.

Bradley MM, Lang PJ. Affective norms for English words (ANEW): Instruction manual and affective ratings. Technical report C-1, the center for research in psychophysiology, University of Florida; 1999 Jan.

Browning H, Veit W. 2022. The sentience shift in animal research. The New Bioethics 28(4) 299–314.

Bush ER, Baker SE, Macdonald DW. 2014. Global trade in exotic pets 2006– 2012. Conservation Biology 28(3) 663–676.

Challender DW, et al. 2021. Mischaracterizing wildlife trade and its impacts may mislead policy processes. Conservation Letters p.e12832.

CITES Secretariat. 2022. World Wildlife Trade Report 2022. Geneva, Switzerland.

CITES and Livelihoods. 2023. https://cites.org/eng/prog/livelihoods accessed 9/7/2023.

Conservation Breeding Specialist Group. 2023. https://www.cpsg.org/our-approach/one-plan-approach-conservation

Dahlstrom MF. 2014. Using narratives and storytelling to communicate science with nonexpert audiences. Proceedings of the National Academy of Sciences 111(4) 13614–13620.

De la Fuente, MF et al. 2023. Keeping reptiles as pets in Brazil: keepers’ motivations and husbandry practices. Journal of Ethnobiology and Ethnomedicine, 19(1), p.46.

Doughnut Economics. 2023. https://doughnuteconomics.org/about-doughnut-economics

Easter T, Trautmann J, Gore M, Carter N. 2023. Media portrayal of the illegal trade in wildlife: The case of turtles in the US and implications for conservation. People and Nature. 10.1002/pan3.10448

Edwards DP, et al. 2021. The dangers of misrepresenting wildlife trade: Response to Natusch et al. (2021). Conservation Biology: the journal of the Society for Conservation Biology. 35(5) 1692-1694.

Entman RM. 1993. Framing: Toward clarification of a fractured paradigm. Journal of Communication. 43(4) 51–58.

Feber RE, Raebel EM, D’Cruze N, Macdonald DW, Baker SE. 2017. Some animals are more equal than others: wild animal welfare in the media. BioScience 67(1) 62–72.

Feinerer I, Hornik K, Meyer D. 2008. Text Mining Infrastructure in R. Journal of Statistical Software 25(5) 1–54. doi:10.18637/jss.v025.i05.

Fitzpatrick LD, Pasmans F, Martel A, Cunningham, AA. 2018. Epidemiological tracing of *Batrachochytrium salamandrivorans* identifies widespread infection and associated mortalities in private amphibian collections. Scientific Reports 8(1) 1–10.

Galtung J, Ruge MH. 1965. The structure of foreign news: The presentation of the Congo, Cuba and Cyprus crises in four Norwegian newspapers. Journal of Peace Research 2(1) 64–90.

Gore ML, Griffin E, Dilkina B, Ferber A, Griffis SE, Keskin BB, Macdonald J. 2023. Advancing interdisciplinary science for disrupting wildlife trafficking networks. Proceedings of the National Academy of Sciences 120(10) p.e2208268120.

Guo X, Meijboom FL. 2023. The development of animal welfare science in China: An explorative analysis. Animal Welfare, 32, p.e72.

Haddon C, Burman OH, Assheton P, Wilkinson A. 2021. Love in Cold Blood: Are Reptile Owners Emotionally Attached to Their Pets?. Anthrozoös 1–11.

Hammond NL, Dickman, A, Biggs, D. 2022. Examining attention given to threats to elephant conservation on social media. Conservation Science and Practice 4(10) p.e12785.

Hughes AC. 2021. Wildlife trade. Current Biology 31(19) R1218–R1224.

IPBES 2019. Global assessment report on biodiversity and ecosystem services of the Intergovernmental Science-Policy Platform on Biodiversity and Ecosystem Services. E. S. Brondizio, J. Settele, S. Díaz, and H. T. Ngo (editors). IPBES secretariat, Bonn, Germany. 1148 pages. 10.5281/zenodo.3831673

Jacobs LR, Page BI. 2005. Who influences US foreign policy? American Political Science Review 99(1) 107–123.

Katz Y. 2013. Against storytelling of scientific results. Nature Methods 10(11) 1045–1045.

Keskin BB, Griffin EC, Prell JO, Dilkina, B, Ferber A, MacDonald J, Hilend R, Griffis S, Gore ML. 2022. Quantitative investigation of wildlife trafficking supply chains: A review. Omega p.102780.

Khelifa R, Amano T, Nuñez MA. 2022. A solution for breaking the language barrier. Trends in Ecology & Evolution 37(2) 109–112.

King TA. 2019. Wild caught ornamental fish: a perspective from the UK ornamental aquatic industry on the sustainability of aquatic organisms and livelihoods. Journal of Fish Biology 94(6) 925–936.

Lockwood JL, et al. 2019. When pets become pests: the role of the exotic pet trade in producing invasive vertebrate animals. Frontiers in Ecology and the Environment 17(6) 323–330.

Mandimbihasina AR, Woolaver LG, Concannon LE, Milner-Gulland EJ, Lewis RE, Terry AM, Filazaha N, Rabetafika LL, Young RP. 2020. The illegal pet trade is driving Madagascar’s ploughshare tortoise to extinction. Oryx 54(2) 188–196.

Marshall BM, Strine C, Hughes AC. 2020. Thousands of reptile species threatened by under-regulated global trade. Nature Communications 11(1) 4738.

Mata F, Araujo J, Soares L, Cerqueira JL. 2023a. Local people standings on existing farm animal welfare legislation in the BRIC countries and the USA. Comparison with Western European Legislation. Journal of Applied Animal Welfare Science, pp.1–14.

Mata F, Dos-Santos M, Cocksedge J. 2023b. Attitudinal and Behavioural Differences towards Farm Animal Welfare among Consumers in the BRIC Countries and the USA. Sustainability, 15(4), p.3619.

Mellor DJ. 2016. Moving beyond the “five freedoms” by updating the “five provisions” and introducing aligned “animal welfare aims”. Animals 6(10) p.59.

Meyer M. 2010. The rise of the knowledge broker. Science Communication 32(1) p.118–127.

Mohammad SM, Turney PD. Crowdsourcing a word–emotion association lexicon. Computational intelligence. 2013 Aug;29(3):436–65. 10.1111/j.1467-8640.2012.00460.x

Moorhouse TP, Balaskas M, D’Cruze NC, Macdonald DW. 2017. Information could reduce consumer demand for exotic pets. Conservation Letters 10(3) 337–345.

Natusch DJ, Lyons JA, Riyanto A, Khadiejah S, Shine R. 2019. Detailed biological data are informative, but robust trends are needed for informing sustainability of wildlife harvesting: A case study of reptile offtake in Southeast Asia. Biological Conservation 233 83–92.

Natusch DJ, Aust PW, Shine R. 2021a. The perils of flawed science in wildlife trade literature. Conservation Biology 35(5) 1396–1404.

Natusch DJ, Aust PW, Shine R. 2021b. Pitfalls in evaluating the sustainability of wildlife trade: reply to Sosnowski and Petrossian and Edwards et al. Conservation Biology 35(5) 1695–1697.

Nijman V, Langgeng A, Birot H, Imron MA, Nekaris KAI. 2018. Wildlife trade, captive breeding and the imminent extinction of a songbird. Global Ecology and Conservation 15 p.e00425.

Obura D, et al. 2023. Prioritizing sustainable use in the Kunming-Montreal global biodiversity framework. PLOS Sustainability and Transformation 2(1) p.e0000041.

Page MJ, McKenzie JE, Bossuyt PM, Boutron I, Hoffmann TC, Mulrow CD, Shamseer L, Tetzlaff JM, Akl EA, Brennan SE, Chou R. 2021. The PRISMA 2020 statement: an updated guideline for reporting systematic reviews. International Journal of Surgery. 88. p.105906.

Pasmans F, Bogaerts S, Braeckman J, Cunningham AA, Hellebuyck T, Griffiths RA, Sparreboom M, Schmidt BR, Martel A. 2017. Future of keeping pet reptiles and amphibians: towards integrating animal welfare, human health and environmental sustainability. Veterinary Record 181(17) 450–450.

Ramírez-Castañeda V. 2020. Disadvantages in preparing and publishing scientific papers caused by the dominance of the English language in science: The case of Colombian researchers in biological sciences. PloS One 15(9) p.e0238372.

R Core Team. 2022. R: A language and environment for statistical computing. R Foundation for Statistical Computing, Vienna, Austria. URL https://www.R-project.org/

Richards SA. Testing ecological theory using the informationLtheoretic approach: examples and cautionary results. Ecology. 2005 Oct;86(10):2805–14. DOI 10.1890/05-0074

Robinson JE, Griffiths RA, Fraser IM, Raharimalala J, Roberts DL, St John FA. 2018a. Supplying the wildlife trade as a livelihood strategy in a biodiversity hotspot. Ecology and Society 23(1).

Robinson JE, Fraser IM, St John FAV, Randrianantoandro JC, Andriantsimanarilafy RR, Razafimanahaka JH, Griffiths RA, Roberts DL. 2018b. Wildlife supply chains in Madagascar from local collection to global export. Biological Conservation 226 144–152.

Ross-Hellauer T, Reichmann S, Cole NL, Fessl A, Klebel T, Pontika N. 2022. Dynamics of cumulative advantage and threats to equity in open science: a scoping review. Royal Society Open Science, 9(1), p.211032.

Santos BS, Crowder LB. 2021. Online news media coverage of sea turtles and their conservation. BioScience 71(3) 305–313.

Scottish Animal Welfare Commission. 2022. https://www.gov.scot/publications/final-report-exotic-pet-working-group-scottish-animal-welfare-commission/

Sengottuvel RR, Mendis A, Sultan N, Shukla S, Chaudhuri A, Mendiratta, U. 2023 From pets to plates: network analysis of trafficking in tortoises and freshwater turtles representing different types of demand. Oryx 1–12 doi:10.1017/S0030605323000376

Shaffer ML, 1981. Minimum population sizes for species conservation. BioScience 31(2) 131–134.

Shiffman DS, et al. 2020. Inaccurate and biased global media coverage underlies public misunderstanding of shark conservation threats and solutions. iScience 23(6) 101205.

Silge J, Robinson, D. 2017. Text mining with R: A tidy approach. JOSS, 1(3). doi:10.21105/joss.00037, 10.21105/joss.00037.

Sinclair JS, Stringham OC, Udell B, Mandrak NE, Leung B, Romagosa CM, Lockwood JL. 2021. The International Vertebrate Pet Trade Network and insights from US imports of exotic pets. BioScience 71(9) 977–990.

Sosnowski MC, Petrossian GA. 2021. Illegal and legal wildlife trade analysis discourse: response to Natusch et al. (2021). Conservation Biology 1 p.3.

Species Use Database. 2023. https://speciesusedatabase.com/ accessed 9/7/2023.

Spooner SL, Walker SL, Dowell S, Moss A. 2023. The value of zoos for species and society: The need for a new model. Biological Conservation 279 109925.

Steinberg A. 2011. Space policy responsiveness: The relationship between public opinion and NASA funding. Space Policy 27(4) 240–246.

Stimson JA, MacKuen MB, Erikson RS. 1995. Dynamic representation. American political science review 89(3) 543–565.

Tingley MW, Harris JBC, Hua F, Wilcove DS, Yong DL. 2017. The pet trade’s role in defaunation. Science 356(6341) 916–916.

Walker JM, Godley BJ, Nuno A. 2019. Media framing of the Cayman Turtle Farm: Implications for conservation conflicts. Journal for Nature Conservation 48 61–70.

Wallack L, DeJong W. 1995. Mass media and public health: Moving the focus from the individual to the environment. Pages 253-268 in Martin SE, Mail PD, eds. The effects of the mass media on the use and abuse of alcohol.

Watters F, Stringham O, Shepherd CR, Cassey P. 2022. The U.S. market for imported wildlife not listed in the CITES multilateral treaty. Conservation Biology 36 p.e13978. 10.1111/cobi.13978

Wickham H. Programming with ggplot2. Ggplot2: elegant graphics for data analysis. 2016:241-53. https://ggplot2.tidyverse.org.

World Health Organisation. 2023. https://www.who.int/europe/initiatives/one-health

