## Supplementary figures and images for "Exploring media representation of the exotic pet trade: taxonomic, framing, and language biases in peer-reviewed publications and newspaper articles"

### Fig S1

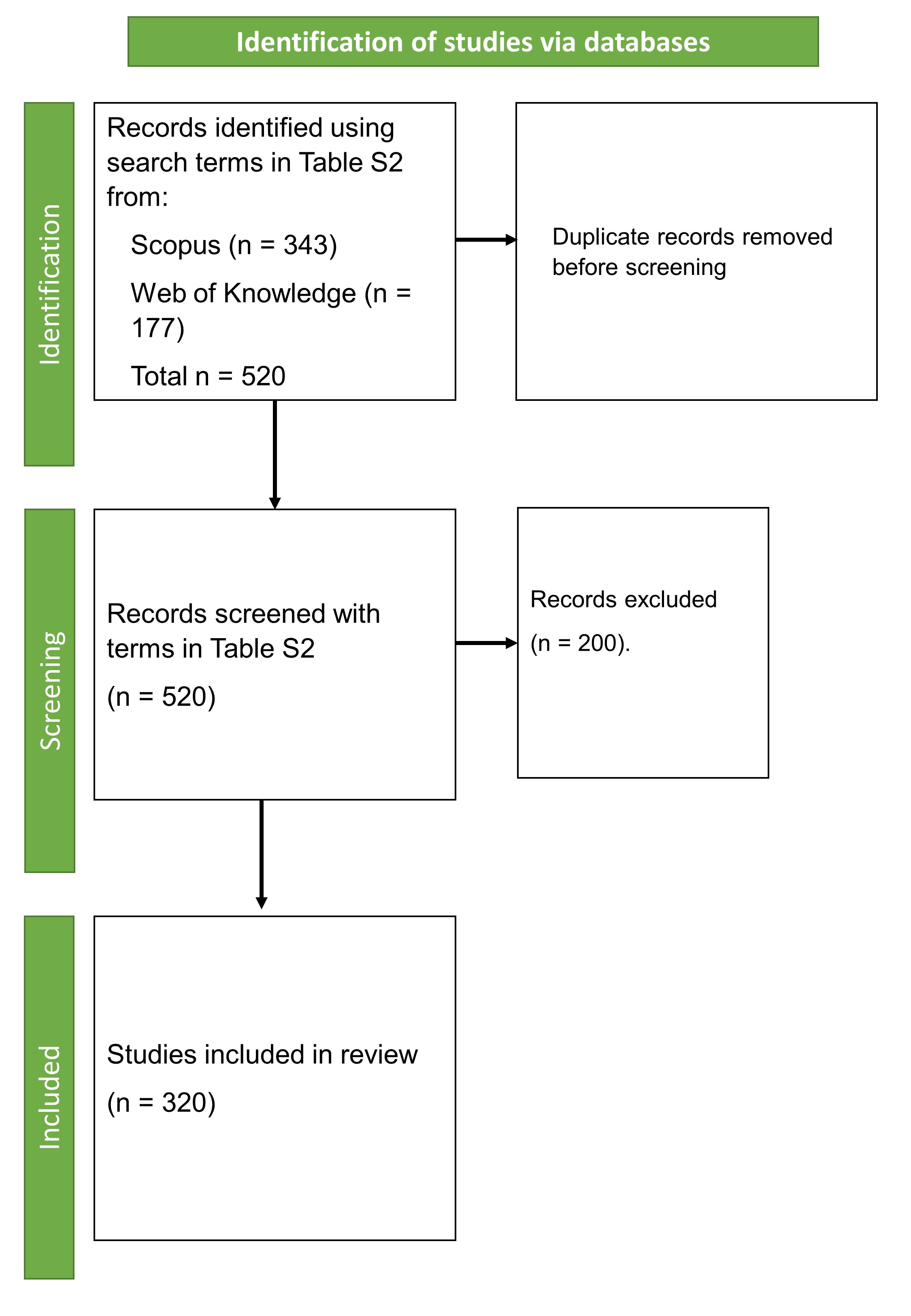
