## Supplementary material for "Exploring media representation of the exotic pet trade: taxonomic, framing, and language biases in peer-reviewed publications and newspaper articles": File S1

**Peer-reviewed literature**

The terms used for searches are presented in S1 Table. The full dataset was filtered for peer-reviewed publications containing one or more of the specific terms in their titles, abstracts, or keywords. For quality control each search was manually screened and documents were excluded if they were not scholarly peer-reviewed publications, mentioned any of the chosen search terms in an irrelevant context, did not mention exotic animals, or only mentioned trade of animals for their parts or as a food or medicine source. To further filter the search results and identify papers whose focus was the exotic pet trade, a list of terms relevant to the pet trade was compiled, based on observations of the documents in our dataset (S2 Table), and the list was then used to filter the dataset for papers containing one or more of those terms in their titles, abstracts or keywords. The search terms were developed to maximise the capture of peer-reviewed publications relating to the exotic pet trade, whilst minimising entries focussed upon the wildlife trade for other reasons (e.g., food). While this approach may have omitted some papers that included trade in animals for multiple reasons (e.g., animals collected for both use as pets and food), for the purposes of this study we used a conservative approach to be more certain that the main patterns in the data were related to animals used for the exotic pet trade rather than predominantly for other purposes. Data cleaning and manipulation were carried out within statistical software R [1], using the following packages: dplyr [2], stringr [3] and tidyverse [4].

**Newspaper articles**

Duplicate articles were removed (e.g, the same article published under a different title, or the same title in a different newspaper). We decided to focus on newspaper articles as a form of popular media because it was more feasible to collect framing data on printed articles rather than other forms of popular media (e.g., television) and because newspaper articles provide a more direct comparison with peer-reviewed papers than would other forms of popular media. We focussed on English language newspaper articles to provide a direct comparison of language use with peer-reviewed publications. The results obtained were uploaded to NVivo [5], which allows qualitative and mixed-methods research, especially the analysis of unstructured text. The software R [1] was used for handling and interrogating text files, and for statistical analysis throughout.

**Sentiment text examples**

Below are examples of a peer-reviewed abstract and a newspaper article. The former obtained a high score for ‘Trust’, an nrc emotion category that was consistently higher in peer-reviewed abstracts than newspaper articles (14.5-17.5% vs, 12.0-12.6%). The latter obtained a high score for ‘Fear’, an nrc emotion category that was consistently higher in newspaper articles than peer-reviewed abstracts (11.4-12.1% Vs. 8.1-11.0%). These example texts have been chosen because they are examples in which one media type (peer-reviewed publication or newspapers) exhibited a higher proportion of a words of that nrc emotion category than the other media type (Table 5 in the main text for details).

Peer-reviewed article abstract text scoring highly for ‘Trust’:

*combating the surge of illegal wildlife trade (iwt) devastating wildlife populations is an urgent global priority for conservation. there are increasing policy commitments to take action at the local community level as part of effective responses. however, there is scarce evidence that in practice such interventions are being pursued and there is scant understanding regarding how they can help. here we set out a conceptual framework to guide efforts to effectively combat iwt through actions at community level. this framework is based on articulating the net costs and benefits involved in supporting conservation versus supporting iwt, and how these incentives are shaped by anti-iwt interventions. using this framework highlights the limitations of an exclusive focus on top-down, enforcement-led responses to iwt. these responses can distract from a range of other approaches that shift incentives for local people toward supporting conservation rather than iwt, as well as in some cases actually decrease the net incentives in favor of wildlife conservation.*

Newspaper article text scoring highly for ‘Fear’:

*At the root of the government's recent scramble to contain the outbreak of monkeypox lies a simple fact: anyone arriving in the United States carrying meat, fruit or a potted plant from any foreign destination is subject to a thorough inspection and confiscation of the item to ensure it isn't harboring diseases or parasites. But an importer of live exotic animals, say Gambian giant pouched rats that are blamed for introducing the monkey pox virus into the United States from Africa and passing it to humans via pet prairie dogs, faces no such check. Gambian rats, and hundreds of other exotic wildlife species, have a far easier time entering the United States than dogs, cats, livestock, horses and people. This latest outbreak of yet another alien disease results from the government's failure to regulate the flow of millions of wild creatures into this country for the pet trade. A veritable Noah's Ark of exotic wildlife carrying viruses, bacteria and parasites that can transmit endemic foreign contagions to humans and to native wildlife are being imported into the United States with scant federal regulation, restriction or precaution. America's craze for exotic pets has created a freewheeling, virtually unregulated wildlife import industry that may account for nearly half of the roughly $30 billion market for pets and pet products in this country. The industry is in serious need of controls. Everything from dangerous carnivores to omnivorous fish to venomous reptiles and amphibians are sold in pet stores, on the Internet, by mail order catalog, at regional auctions and in local swap meets. Animals have long been known to transmit zoonotic illnesses to humans. They include E. coli, rabies, salmonella, trichinosis, yellow fever, malaria, botulism, streptococcus and influenza. What are known as "emerging diseases" recently have increasingly jumped from animals to humans as contact with exotic creatures has risen and opportunistic infectious agents have found new hosts. They include HIV/AIDS, Hepatitis B, the hemorrhagic Ebola and Marburg viruses, Lyme disease, hantavirus, mad cow disease, West Nile virus, the respiratory killer SARS and now monkeypox. Experts believe this animal-human crossover could spawn dangerous new pathogens and increase the chances for another deadly disease outbreak. The Humane Society of the United States began campaigning against exotic animal imports 30 years ago when it supported a successful petition to the U.S. Food and Drug Administration to ban the import and sale of small turtles that carry salmonella. In 1975, the government banned imports of all primates for the pet trade because they carry several dangerous diseases. Following the monkeypox outbreak, the government banned the import, sale and distribution of Gambian rats and other African rodents and halted trade in native American prairie dogs. The government's practice of targeting wildlife after a disease outbreak illustrates a major flaw in public health protection -- closing the barn door after the horse has bolted. According to the Centers for Disease Control and Prevention, there are 9 million pet reptiles -- snakes, iguanas, lizards and turtles -- in the United States, and they are responsible for about 90,000 cases of salmonella poisoning annually. The disease causes severe diarrhea, fever, vomiting, even death -- with children and the elderly the most vulnerable. Four years ago the Humane Society petitioned the FDA for an import ban on all pet reptiles in response to the soaring incidence of salmonellosis. We are still awaiting the agency's response. Government defenses against the exotic animal disease threat are fragmented among several federal agencies that regulate imports of dogs, cats, livestock, horses, meat and produce. Everything else gets waved through. Says an Agriculture Department spokesman: "We don't regulate importation of fish, reptiles, lions, tigers, bears, foxes, monkeys, endangered species, guinea pigs, hamsters, gerbils, mice, rats, chinchillas, squirrels, mongooses, chipmunks, ferrets and other rodents." Exotics can also wreak ecological and financial havoc by introducing diseases to domestic wildlife, livestock, poultry and fish populations which have no natural resistance to them. A recent case: Exotic Newcastle Disease, carried into California this year by smuggled Mexican parakeets and initially spread to four other states by illegal cock fighters whose game fowl became infected. The disease has forced the government to destroy 3.5 million chickens and turkeys and has cost taxpayers more than $180 million. When millions of surplus cats and dogs are euthanized every year because homes cannot be found for them, there is no good reason to take wild animals from their natural habitats and confine them to a tiny cage or a small enclosure for the rest of their lives. Consumers should consider the health risks and the humane issues associated with any species of wild animal -- exotic or native -- obtained as a pet. Any time a wild creature is brought into the home, it can bring with it every bacteria, virus or parasite it has been exposed to. Even with a lengthy quarantine, there is no way to assure that these animals are healthy or will not pass on disease-causing pathogens to humans. Until a sound system to protect public health is in place, the federal government should prohibit imports of all exotic mammals, reptiles, amphibians and birds -- wild caught or captive bred -- destined for the pet trade. If someone wants a loving pet, there are plenty available for adoption at local humane societies and breed rescue groups.*

**Example of narrative-driven newspaper article versus data-driven peer-reviewed abstract.**

**Narrative-driven newspaper article**

“Exotic pet trade helped spread devastating fungus, study finds”

Invasive fungus probably spread via international trade, researchers say Professor Matthew Fisher went deep into the gloomy rain forest of French Guiana to catch poison dart frogs on behalf of science. It was slippery, soggy, vaguely reptilian work. Fisher, whose work uniform was a pair of shorts, discovered that the best way to capture a frog was by slithering. "You've got to pretend you're a snake," said the epidemiologist from Imperial College London. He would silently creep to within an arm's length of his target and then lunge forward: "You've just got to clap both of your hands around it and hold on tight." The frogs were critical elements of a 10-year global investigation by 38 research institutions of a pathogenic fungus, Batrachochytrium dendrobatidis (Bd), that is devastating amphibian populations around the world. The fungus, called a chytrid, causes an often-fatal skin disease, chytridiomycosis. Fisher and his colleagues cultured samples of the fungus, ran genetic tests and sought to understand when and where the pathogen emerged and how it spread around the planet. Their results, published Thursday in the journal Science, indicate that the Bd fungus pandemic did not begin 23,000 years ago, as one earlier hypothesis suggested, but rather sometime in the 1900s. Global trade and the marketing of exotic pets probably propelled it. The genetic signals point to a common ancestor in East Asia, possibly on the Korean Peninsula. Fisher said the surge in activity in East Asia during World War II and the Korean War, and the increased movement of people and cargo, could have played a role in distributing fungus-infected frogs and toads to other parts of the world. "When you globalize trade, you globalize unexpected secondary consequences of trade," Fisher said. Karen Lips, a University of Maryland biologist who has documented declines in amphibian populations in Central America, said the report shows that the chytrid fungi are more widespread and diversified than previously known. Such fungi are an invasive species and have a devastating impact on biodiversity, yet they don't get the kind of attention given to more charismatic alien species - such as Burmese pythons, brown tree snakes, Northern snakeheads, feral hogs and rabbits. Lips, who was not involved in the new study, said it shows there are many lineages of fungus, and "these things can hybridize. . . . The impacts are global and much larger than we've seen before." Not just frogs and toads are threatened by the chytrid fungi. Salamanders in Europe have been slammed by such a fungus, Batrachochytrium salamandrivorans (Bsal), that emerged in Vietnam. Its arrival in the United States would imperil this country's salamander population. It was in the 1980s that field researchers noted a global decline in amphibian populations. No one could explain it. One hypothesis pointed to the thinning ozone layer. Only in the 1990s did the fungus explanation surface. Some frog species have already gone extinct in the wild, including the mountain chicken frog, said Simon O'Hanlon, a research associate at Imperial College London and the lead author of the new report. There are roughly 7,800 species of frogs and toads globally, and only about 1,400 have been tested for the fungus. About half showed signs of infection. Frogs and toads are marketed globally as exotic pets, a food source and for use in scientific research. They can also be stowaways. Asian toads, for example, recently invaded Madagascar by hiding in imported mining equipment. The new report notes that it's probably not a coincidence that the estimated dates for the emergence of the chytrid pandemic are roughly the same as for the "big bang" in international trade. "We're shipping whatever around the world with no regulation, moving things from one environmental niche to another," O'Hanlon said. "It seems that we're putting our global biodiversity recklessly at risk." The researchers said they need more data, meaning more captured frogs and toads from which to glean genetic information about the pandemic fungus. Fisher ultimately nabbed 600 poison dart frogs in French Guiana, taking small tissue samples before releasing them back into the rain forest. He stressed the importance of wearing gloves in this work. "These frogs are absolutely lethal," he said. "If you lick them, it's curtains.”

**Data-driven peer-reviewed abstract**

Wombwell, E.L., Garner, T.W., Cunningham, A.A., Quest, R., Pritchard, S., Rowcliffe, J.M. and Griffiths, R.A., 2016. Detection of Batrachochytrium dendrobatidis in amphibians imported into the UK for the pet trade. *EcoHealth*, *13*, pp.456-466.

“*There is increasing evidence that the global spread of the fungal pathogen Batrachochytrium dendrobatidis (Bd) has been facilitated by the international trade in amphibians. Bd was first detected in the UK in 2004, and has since been detected in multiple wild amphibian populations. Most amphibians imported into the UK for the pet trade from outside the European Union enter the country via Heathrow Animal Reception Centre (HARC), where Bd-positive animals have been previously detected. Data on the volume, diversity and origin of imported amphibians were collected for 59 consignments arriving at HARC between November 2009 and June 2012, along with a surveillance study to investigate the prevalence of Bd in these animals. Forty-three amphibian genera were recorded, originating from 12 countries. It was estimated that 5000–7000 amphibians are imported through HARC into the UK annually for the pet trade. Bd was detected in consignments from the USA and Tanzania, in six genera, resulting in an overall prevalence of 3.6%. This suggests that imported amphibians are a source of Bd within the international pet trade*.”

**References**

[1] R Core Team, 2021. R: A language and environment for statistical computing. R Foundation for Statistical Computing, Vienna, Austria. URL <https://www.R-project.org/>

[2] Wickham H, François R, Henry L, Müller K, Vaughan D. 2023. dplyr: A Grammar of Data Manipulation. https://dplyr.tidyverse.org, https://github.com/tidyverse/dplyr.

[3] Wickham H. 2022. stringr: Simple, Consistent Wrappers for Common String Operations. https://stringr.tidyverse.org, https://github.com/tidyverse/stringr.

[4] Wickham H, et al. 2019. Welcome to the tidyverse. *Journal of Open Source Software*, **4**(43), 1686, https://doi.org/10.21105/joss.01686

[5] QSR International Pty Ltd. 2018. NVivo (version 12). Retrieved from

<https://www.qsrinternational.com/nvivo-qualitative-data-analysis-software/home>
