## Supplementary material for "Exploring media representation of the exotic pet trade: taxonomic, framing, and language biases in peer-reviewed publications and newspaper articles": Table S1

S1 Table: Details of terms used for peer-reviewed scientific literature search. Note: words containing * allow for variants to appear in search results, including pluralisation.

| Database | Document Section | Boolean Operator | Search Terms |
| --- | --- | --- | --- |
| Scopus | Title, Abstract & Keywords |  | exotic* pet* OR wildlife trade |
| Scopus | Title, Abstract & Keywords | AND NOT | “wet market” or food or agriculture or livestock or cattle or rhinoceros or elephant |
| Web of Science | Title |  | exotic* pet* OR wildlife trade |
| Web of Science | Topic | NOT | “wet market” or food or agriculture or livestock or cattle or rhinoceros or elephant |
