## Supplementary material for "Exploring media representation of the exotic pet trade: taxonomic, framing, and language biases in peer-reviewed publications and newspaper articles": Table S2

S2 Table: Details of ‘pet trade’ and related terms for secondary screening of articles to ensure a focus on the exotic pet trade. Note: some words are incomplete to allow detection of variants in R, e.g., “aquari” could return “aquarium” or “aquaria”.

| Key Term | Related Terms |
| --- | --- |
| Pet Trade | pet, collect, ornament, poach, captive, aquaculture, aquari*, aviculture, companion, pond stocking, trade, import, sale, translocate, commercial, market |
