## Supplementary material for "Exploring media representation of the exotic pet trade: taxonomic, framing, and language biases in peer-reviewed publications and newspaper articles": Table S3

S3 Table: Details of each framing category and related terms. Note: some words are incomplete to allow detection of variants in R, e.g., “financ” could return “finance” or “financially”.

| Framing Category | Related Terms |
| --- | --- |
| Conservation | conservation, endanger, threat, ecosystem, ecology, overexploit, decline, biodiversity |
| Economy | econom, livelihood, subsistence, income, poverty, developing countr, developing region, financ |
| Disease | disease, spillover, pathog, disease, parasite, infect, bacteria, zoono, pathogen, virus, venom, microb, poison, public health, rabies, tuburculosis, salmonella, allerg, infest, epidem, transmission, toxin, biosecurity |
| Invasive Species | invas, alien, pest, introduce, nonindigenous, release, establish, hybrid, home range |
| Welfare | welfare, husbandry, pain, suffer, enrichment, cage size, stress, veterinary, comfort, injur, pet medicine |
