## Supplementary material for "Exploring media representation of the exotic pet trade: taxonomic, framing, and language biases in peer-reviewed publications and newspaper articles": Table S4

S4 Table: Details of taxonomic foci and related terms. Note: Fish and Crustacean have been combined into a single category, as they are often traded together.

| Taxonomic Group | Related Terms |
| --- | --- |
| Amphibian | amphibian, frog, toad, salamander, newt, axolotl |
| Bird | bird, parrot, passerine, bird of prey, owl, macaw, cockatoo, cockatiel |
| Fish & Crustacean | fish, aquari, cichlid, tetra, shrimp, blue tang, clownfish, betta, crayfish |
| Mammal | mammal, primate, monkey, macaque, ferret, sugar glider, prairie dog, chinchilla, big cat, lion, tiger, ocelot, otter, degu, guinea pig, loris, marmoset, fox, sloth |
| Reptile | reptile, lizard, turtle, terrapin, tortoise, snake, dragon, gecko, chameleon, herptile, crocodil, tuatara, caiman, iguana, skink, cobra, viper, constrictor |
