## Supplementary material for "Exploring media representation of the exotic pet trade: taxonomic, framing, and language biases in peer-reviewed publications and newspaper articles": Table S5

S5 Table: Raw count of peer-reviewed papers in each framing category and combinations thereof.

| Framing | Count of papers (2001-2021) |
| --- | --- |
| Conservation | 61 |
| Disease | 25 |
| Economics | 2 |
| Invasive species | 24 |
| Welfare | 10 |
| No/other frame | 21 |
| Multiple framing | 177 (breakdown below) |
| *Conservation & Invasives* | *29* |
| *Conservation & Economics* | *21* |
| *Disease & Welfare* | *10* |
| *Disease & Invasives* | *9* |
| *Conservation & Welfare* | *9* |
| *Conservation & Disease* | *7* |
| *Economics & Invasion* | *5* |
| *Disease & Economics* | *3* |
| *Invasives & Welfare* | *3* |
| *Economics & Welfare* | *1* |
| *Three or more framings* | *80* |
