## Supplementary material for "Exploring media representation of the exotic pet trade: taxonomic, framing, and language biases in peer-reviewed publications and newspaper articles": Table S6

S6 Table: Example quotations for a selection of framings and nrc emotion categories from peer-reviewed publication abstracts and newspaper articles. Examples of framing categories were selected on the basis of the frequency with which those categories were used and their centrality to the analyses within this overview. Examples of emotion categories were selected on the basis of their frequency of use and differences between media types.

|  | Peer-reviewed publication abstract |  | Newspaper article |
| --- | --- | --- | --- |
| Framing category  Multiple frames | “Abandonment of pets creates numerous negative externalities and multimillion-dollar costs, in addition to severe consequences and problems concerning animal welfare (e.g., starvation, untreated disease, climatic extremes, uncertainty of rescue and adoption), ecological (e.g., invasive species and introduction of novel pathogens), public health and safety (e.g., risks to people from bites, zoonoses, or road hazards), and economic (e.g., financial burdens for governmental and non-governmental organizations).”  From “Amelioration of pet overpopulation and abandonment using control of breeding and sale, and compulsory owner liability insurance” |  |  |
|  |  | **Framing category**  Laws and regulations | “Own a sea turtle, a marmoset or any other exotic species? You'll now have to submit a form to the chief wildlife warden in your state. In a first of its kind move, the Union ministry of environment, forest and climate change has issued an advisory to streamline the process of importing and possessing live exotic animals while also asking owners of exotic species to voluntarily disclose information on their pets within six months.”  From “Govt moves to regulate import of exotic animals [Times Nation]” |
| Framing category  Conservation | “Wildlife trade is the very heart of biodiversity conservation and sustainable development providing an income for some of the least economically affluent people and it generates considerable revenue nationally. In Asia the unsustainable trade in wildlife has been identified as one of the main conservation challenges”  From “An overview of international wildlife trade from Southeast Asia” | **Framing category**  Conservation | "Our wildlife inspectors and special agents work with CBP officials to combat wildlife trafficking at U.S. ports of entry," said Edward Grace, Assistant Director of the U.S. Fish and Wildlife Service Office of Law Enforcement. "The armadillo girdled lizard is a fascinating reptile with distinct natural "armor" that helps it survive in the wild. Unfortunately, one of the biggest threats to this species is the illegal wildlife trade. We would like to thank our partners at CBP and the Cincinnati Zoo and Botanical Garden for their assistance with this case. Together, we can help protect imperiled species throughout the world."  From “Cincinnati customs and border protection rescues lizards from illegal pet trade” |
| Framing category  Welfare | “The wildlife trade threatens global biodiversity and animal welfare, where parrots are among the taxa most frequently traded, supplying exotic pets and captive breeders worldwide.”  From “China's online parrot trade: generation length and body mass determine sales volume via price” | **Framing category**  Welfare | “HUNDREDS of ball python snakes are suffering unnecessary cruelty in the UK after being crammed into small plastic boxes and put up for sale, an animal welfare organisation claims. aid World Animal Protection said the shy and nocturnal animals, which are popular exotic pets in the UK, were being denied the space to move properly, as well as being subjected to bright lights and loud noises. '' These are wild that feel and Investigators for the charity found ball pythons at the meetings, held at Doncaster Racecourse, in "tiny" boxes. It is now urging the venue - home to one of the last reptile markets in the UK - to stop hosting the events.”  From “Let's end cruel market trade of exotic pets” |
| nrc emotion category  Trust | “Nature has the potential to provide wide-ranging economic contributions to society – from ecosystem services to providing income to communities via fair trade of resources.”  From: “Cites and beyond: illuminating 20 years of global, legal wildlife trade” |  |  |
|  |  | **nrc emotion category**  Fear | “This latest outbreak of yet another alien disease results from the government's failure to regulate the flow of millions of wild creatures into this country for the pet trade. A veritable Noah's Ark of exotic wildlife carrying viruses, bacteria and parasites that can transmit endemic foreign contagions to humans and to native wildlife are being imported into the United States with scant federal regulation, restriction or precaution”  From: “Exotic pets need controls” |
|  |  | **nrc emotion category**  Sadness | “Most are unaware of, or are indifferent to, the suffering of wildlife that are traded as pets. Local celebrities and political personalities are among those who have purchased protected species as pets. Comments from social media users are largely encouraging and full of envy, with many expressing the desire to purchase similar animals due to their beauty and the fact that they are seen as status symbols. Reasons wildlife traders and owners give to justify keeping exotic pets include: THE animals' natural habitats have been destroyed, or their mothers have been killed, and the young have no way of surviving in the wild”  From: “Cruelty abounds in exotic pet trade” |
|  |  | **nrc emotion category**  Anticipation | “An MP has called on the Government to review laws regarding the importing of exotic pets in the wake of the coronavirus outbreak. Mark Pritchard, MP for The Wrekin, was commenting on research by the University of California which found close contact with wild animals through hunting, trade or habitat loss put the world at increased risk of new diseases.”  From: “MP calls for curbs on trade in exotic pets” |
